# *Trypanosoma brucei* colonises the tsetse gut via an immature peritrophic matrix in the proventriculus

**DOI:** 10.1101/513689

**Authors:** Clair Rose, Naomi A. Dyer, Aitor Casas-Sanchez, Alison J. Beckett, Carla Solórzano, Ben Middlehurst, Marco Marcello, Michael J. Lehane, Ian A. Prior, Álvaro Acosta-Serrano

## Abstract

The peritrophic matrix (PM) of haematophagus insects is a chitinous structure that surrounds the bloodmeal, forming a protective barrier against oral pathogens and abrasive particles. To establish an infection in the tsetse midgut, *Trypanosoma brucei* must colonise the ectoperitrophic space (ES), located between the PM and gut epithelium. Although unproven, it is generally accepted that trypanosomes reach the ES by directly penetrating the PM in the anterior midgut. Here we revisited this event by employing novel fluorescence and electron microscopy methodologies and found that instead, trypanosomes reach the ES via the newly secreted PM in the tsetse proventriculus. Within this model, parasites colonising the proventriculus can either migrate to the ES or become trapped within PM layers forming cysts that move along the entire gut as the PM gets remodelled. Early proventricular colonisation appears to be promoted by unidentified factors in trypanosome-infected blood, resulting in higher salivary gland infections and potentially increasing parasite transmission.

## Introduction

*Trypanosoma brucei* sub-species, the causative agent of human sleeping sickness and also partially responsible for animal trypanosomiasis in sub-Saharan Africa, are transmitted exclusively by flies of the family Glossinidae, commonly known as tsetse. These parasites have a complex life cycle within the fly, but key to transmission is the ability to first establish an infection within the insect midgut. After a fly ingests blood from an infected mammal, the “stumpy” bloodstream trypanosome transforms into the procyclic stage within the midgut lumen [1]. During this process, the coat of variant surface glycoproteins (VSG) is replaced by a different one composed of procyclins [2, 3]. In the most accepted model of parasite migration within the tsetse, procyclic trypanosomes first establish an infection in the ectoperitrophic space (ES) (defined as the space between the gut epithelium and the peritrophic matrix (PM) Fig. 1a), followed by colonisation of the proventriculus (also known as cardia), and terminating in the salivary glands, where the parasites become mammalian infective again [4–6].

**Fig. 1.**
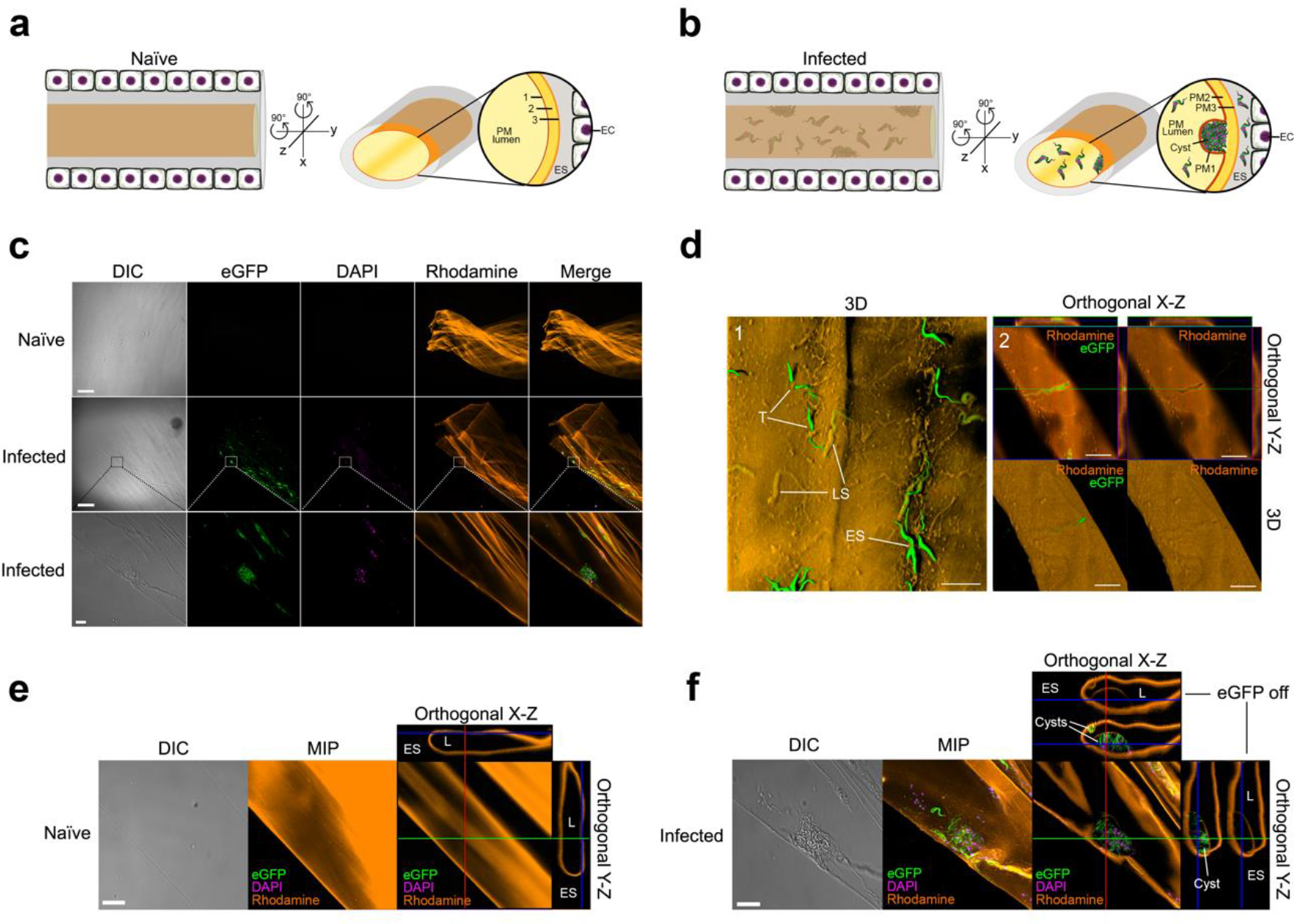
CLSM reveals trypanosome cysts formed between PM layers. **a**, Cartoon depicting a 2D view of naïve tsetse midguts. Rotating 90° on both the X and Y axis gives an indication of what can be seen under CLSM and provides a guide for understanding the orientation of the *ex vivo* PMs in the orthogonal view. **b**, Same as Fig. 1a but depicting a section of a trypanosome-infected gut. Although the three PM layers cannot be seen under CLSM, this schematic shows the position of the trypanosome cysts between PM1 and PM2; these cysts always orientate towards the luminal side of the gut (see also Fig. 2). ES, ectoperitrophic space. EC, epithelial cell. **c**, Washed, *ex vivo* PM from a naïve fly (11 dpi) stained with WGA (top). In infected flies (11 dpi), eGFP-trypanosomes can be seen (green) in close proximity to the PM, with DAPI (magenta) showing parasite nuclei and kinetoplasts (middle). Scale bar 200µm. Inset corresponds to the higher magnification of the same area as seen in bottom panel. DIC, Differential Interference Contrast. Scale bar 20μm. **d**, (1) CLSM 3D reconstructions from multiple z-stacks of washed *ex vivo* PMs from a fly at 9 dpi. Ectoperitrophic space side (ES), luminal side (LS), trapped trypanosomes (T). Scale bar 20 μm. (2) Maximum Intensity Projection (MIP) (top) and 3D reconstruction (bottom) of a trapped trypanosome. **e**, A PM sample from a naïve fly depicting how this tissue looks under DIC and MIP after rendering from multiple z-sections, whilst the orthogonal view shows the XZ/YZ planes of the folded PM section. **f**, DIC and MIP of an infected PM sample containing trypanosome cysts, whilst the orthogonal XZ-YZ views show trypanosomes trapped within PM layers. A second, smaller cyst-like structure can be seen in the XZ orthogonal view at a North-West position to the bigger cyst. ES, ectoperitrophic space. L, lumen. Scale bar 20μm.

The tsetse PM functions to compartmentalise the bloodmeal and to prevent both abrasion and infection of the gut epithelium [7], thus acting as a protective barrier that trypanosomes must overcome in order to reach the ES. *Glossina morsitans* secretes a type II PM, which is continuously produced at a rate of approximately 1 mm/h [8, 9] as an unbroken, multi-layered concentric sleeve (becoming fully formed after ~80-90h of being secreted [10]) by specialised cells in the proventriculus. This immunologically important organ [11], marks the border between the ectodermal foregut (i.e. buccal cavity, pharynx, oesophagus and crop) and entodermal midgut, functioning as a valve due to its arrangement into a ring-shaped fold (valvular cardiaca) [12] (see also Supplementary Fig. 1). After secretion, the tsetse PM is assembled as a trilaminate sheath (PM1-3) [13] (Supplementary Fig. 2), with each layer differing in thickness and composition, but mainly comprised of chitin fibres that are cross-linked to structural glycoproteins (peritrophins) [13–15].

Several suggestions have been made for how trypanosomes reach the tsetse ES, including circumnavigation of the PM in the posterior gut [16, 17], penetration of the ‘freshly secreted’ PM within the proventriculus [18–21], or direct penetration of the ‘mature’ PM within the anterior midgut [22–25]. The latter hypothesis, which involves 1) parasite recognition to, and penetration of, PM1 layer (which faces the gut lumen; Fig. 1a), 2) direct crossing of PM2 and PM3 layers, and 3) exit to the ES, has persisted for over 40 years and has been influenced mainly by the visualisation of ‘penetrating’ trypanosomes between PM layers [22]. However, neither an adhesion ligand on PM1 has been identified nor has experimental evidence for steps 2 and 3 been obtained. Moreover, unlike parasites such as *Leishmania* [26] and *Plasmodium* [27], trypanosomes do not secrete PM-degrading enzymes such as chitinases [28]. Overall, this suggests that the fast turnover of a structurally complex tsetse PM would make a difficult barrier for trypanosomes to degrade, although, very little is known about the physiological response of type II PMs to oral pathogens [29, 30].

In this work, we have revisited how *T. brucei* reaches the tsetse ES by employing several microscopy techniques, including serial block-face scanning electron microscopy (SBF-SEM), and novel confocal laser scanning microscopy (CLSM) methodologies, which collectively allowed the 3D-reconstruction of trypanosome-infected tsetse tissues. We propose that ES invasion occurs via the proventriculus during PM assembly rather than by direct crossing of the mature PM in the midgut, as previously suggested [22, 24, 25]. Furthermore, we give evidence that an early proventricular invasion by trypanosomes is promoted by unknown factor(s) present in trypanosome-infected blood, thus leading to a higher prevalence of salivary gland infections and potentially increasing parasite transmission.

## Results and Discussion

### CLSM shows trypanosomes are trapped within the tsetse PM

In order to visualise how trypanosomes interact with the tsetse PM, we analysed by CLSM *ex vivo* PMs stained with rhodamine-conjugated wheat germ agglutinin (WGA) [31] from either naïve flies or flies infected with eGFP-expressing trypanosomes (*n*=35) (Fig. 1 and Supplementary Videos 1 and 2). WGA exclusively recognises the PM chitin fibres as shown by its inhibition with chitin hydrolysate or when tissues were stained with the succinylated lectin (not shown). Whilst individual trypanosomes appear to be partially penetrating the PM or stuck on either the ES or the luminal side (Fig. 1d), z-stacks orthogonal projections depicted many parasites inside PM cysts as the rhodamine signal could be seen above and below the cells (Fig. 1f). This is better visualised when the eGFP signal is switched off. Moreover, the integrity of all PM cysts analysed was never compromised (i.e. no evidence of parasites penetrating any of the PM layers) and their thinner part (i.e. PM1, see EM section) always faced the luminal side.

### Transmission electron microscopy (TEM) analyses of trypanosome-infected midguts

TEM was then used to better understand, at the ultrastructural level, the nature of the PM cysts and the overall localisation of parasites in infected midguts. We initially focused on the anterior midgut as previous work suggested trypanosomes may cross the PM in this region [22–25]. Parasites were observed either in the lumen, trapped within PM layers or already in the ES at all time-points (5, 8 and 11 dpi). In most infected flies, PM damage was a common occurrence, which was typically characterised by a separation of PM1 and PM2 layers (Fig. 2a-c, e, and f), as previously reported [22–24]. PM1 appears as a thin (electron-dense) layer that is equivalent to the luminal rhodamine signal observed by CLSM (Fig. 1f). Furthermore, this damage was not observed in naïve or refractory flies (not shown), and usually one or more trypanosomes were found within this separation. Occasionally, parasites were seen embedded within PM2 rather than ‘unzipping’ PM1 from PM2 (Fig. 2d). Moreover, at 11 dpi, multiple parasites were commonly found between PM layers, forming bigger cysts (Fig. 2f). The presence of trypanosome-filled cysts, which were more commonly observed in older infected flies, agrees with the structures observed by CLSM (Fig. 1b and d) and previous observations [25, 32]. Interestingly, in all instances where PM1 separated from PM2, we found no evidence of breaks, degradation or thinning of PM1, even when cysts appear to contain high parasite numbers (Fig. 2f). Furthermore, we never observed parasites in the process of entering or leaving the PM1 or PM2 side, partially in or out of the PM, nor did we see a complete break through PM2 layer.

**Fig. 2.**
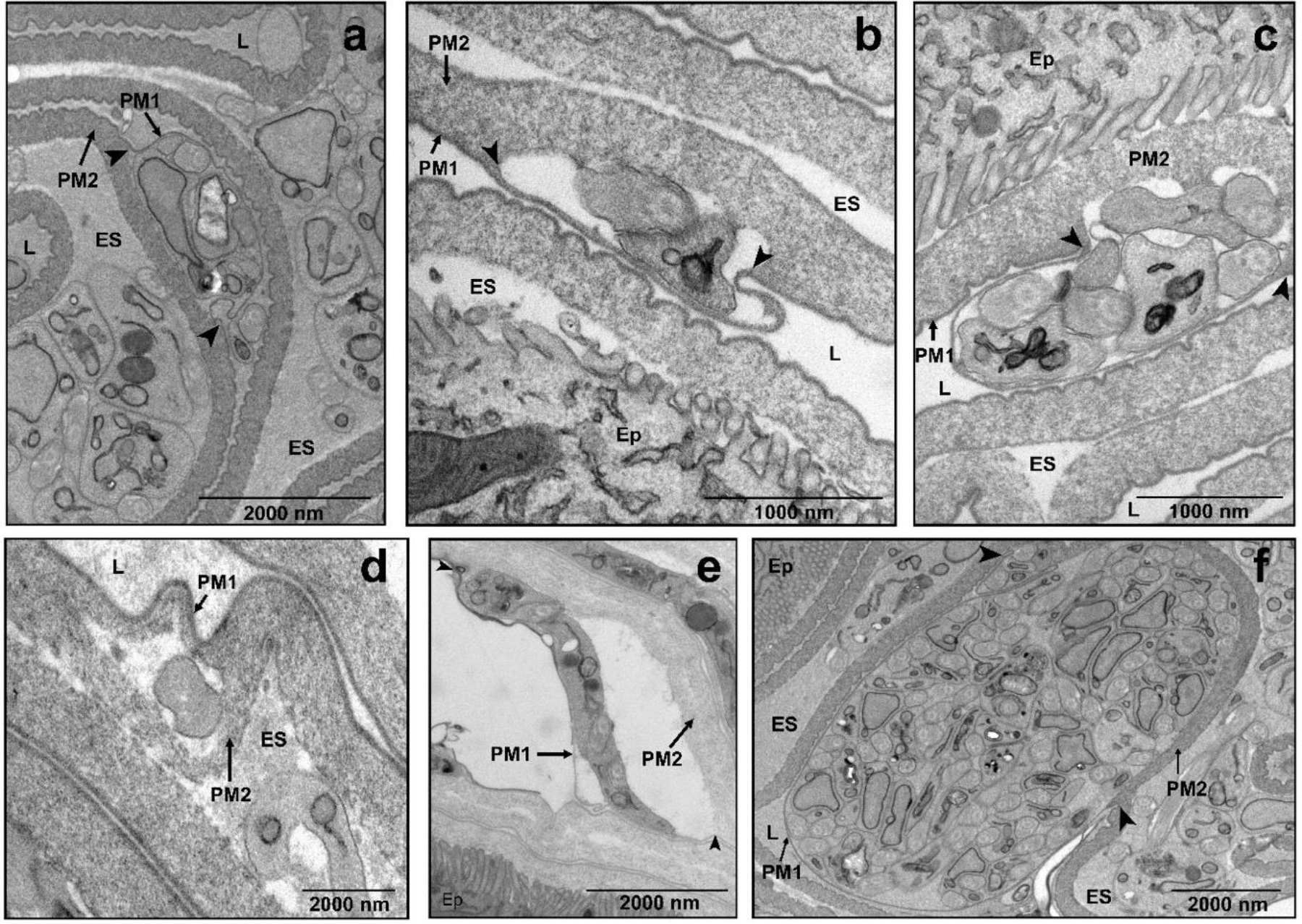
TEM images of sections showing trypanosomes between tsetse PM layers. Typical damage found in infected flies is the separation of PM1 from PM2. Usually, the electron dense (PM1) layer appears unbroken, but can be seen peeling away from the second layer (arrowheads). Note that PM1 remains unbroken even when cysts contain high parasite numbers (**f**). Images were taken from flies at 11 dpi (**a**-**c**, and **f**), 8 dpi (**d**) or 5 dpi (**e**). L, lumen. ES, ectoperitrophic space. Ep, epithelial cells. Numbers of technical and biological replicates used, average number of grids and average number of images per separate grid can be seen in Supplementary table 1.

### SBF-SEM analysis of a trypanosome cyst reveals conserved parasite orientation and absence of PM degradation

To gain more insights into the organisation of trypanosomes within PM cysts located in the anterior midgut, we used SBF-SEM. We prepared >500 serial sections (each ~100nm thick) of a cyst sample (from a fly at 11 dpi) and then 3D-reconstructed this region (Fig. 3, Supplementary Video 3 and 4). It was observed that all parasites, which appeared to be aligned in the same direction as indicated by the orientation of the flagellar tips, were exclusively contained within PM1 and PM2 (Fig. 3c). However, no evidence of crossing or PM damaged induced by trypanosomes was seen corroborating the CLSM and TEM observations (Fig. 1 and Fig. 2).

**Fig. 3.**
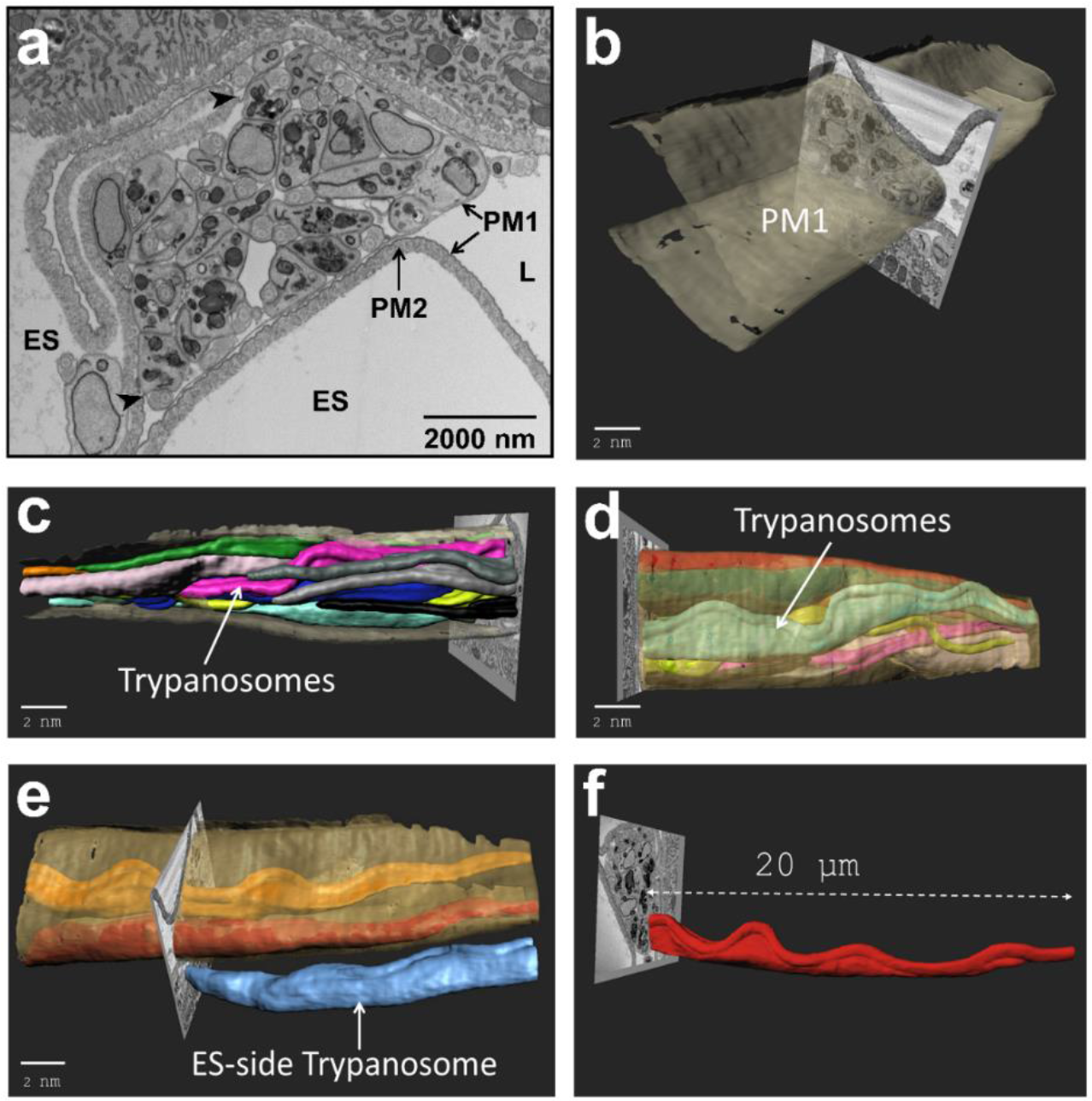
SBF-SEM 3D reconstruction of a trypanosome cyst in the PM from the anterior midgut. Sample taken from a fly at 11 dpi. See also Supplementary Video 4. **a**, Multiple trypanosomes between PM1 and PM2. Arrows show the point of separation as trypanosomes reside inside and both layers remain unbroken. ES, Ectoperitrophic Space. L, Lumen. **b**-**e**, SBF-SEM slices merged with manual segmentation. **b**, Image illustrating breaks or damage to PM1 (grey) are absent during a trypanosome infection. **c**, Multiple parasites between PM1 and PM2; most parasites appeared oriented in the same direction as indicated by the position of the anterior end flagellar tips. **d**, A reverse view of the image depicted in 4c, showing parasites contained within PM1. **e**, Still depicting one parasite (blue) in the ectoperitrophic space side. **f**, A measurement of a partially reconstructed trypanosome within the cyst. Scale bars are representative of the SEM image (not the reconstruction).

Why most trypanosomes trapped within the cyst (Fig. 3) appear to have the same orientation is unknown, particularly when there is no evidence of cell duplication (e.g. flagellar division) in this and other cysts that were analysed by TEM. Alternatively, we hypothesise that proventricular parasites may form cysts by collective motion (CoMo) [31] whereby several trypanosomes, swimming in the same direction, may simultaneously penetrate through an immature PM thus becoming trapped between its layers.

### Trypanosomes reach the ES by early invasion of the proventriculus

The fact that none of the trypanosomes inside the reconstructed cyst or the ones observed by either TEM or CLSM appear to penetrate the PM layers was puzzling. This raises the question of how these cysts are formed if no evidence of parasite crossing is seen in the anterior midgut. One clue, however, came from the lengths of individual trypanosomes from inside the (3D-reconstructed) cyst, which were longer than average midgut procyclics (~20μm; see example in Fig. 3f) and so similar in size to mesocyclic proventricular forms [31, 33] (see also Fig. 7). Therefore, we hypothesised that trypanosome-containing cysts could originate in the proventriculus during PM assembly and consequently, analysed the proventriculus from infected flies at 5 (early infection, Fig. 4) and 11 (late infection, Fig. 5) dpi. After 5 dpi, the proventriculus was heavily infected (63.6% prevalence) with trypanosomes (Fig. 4b-e). Parasites were observed to be adjacent to where the foregut cells become confluent with midgut cells. This suggests trypanosomes can overcome or bypass the PM at this point (Fig. 4b), confirming previous observations [18–21]. Parasites were also observed in the lumen and ectoperitrophic side of the PM and, in some cases, in close proximity to (but not penetrating) the epithelial cells. Moreover, they could also be seen between PM1 and PM2 (Fig. 4d). A proventriculus from a fly at 5 dpi (Supplementary Fig. 3) was subsequently processed for SBF-SEM at the regions of interest (ROI) shown (Supplementary Video 5 and 6), and a partial reconstruction of a small number of parasites that were in close proximity to the chitinous foregut was performed on ROI 2 (Supplementary Video 7).

**Fig. 4.**
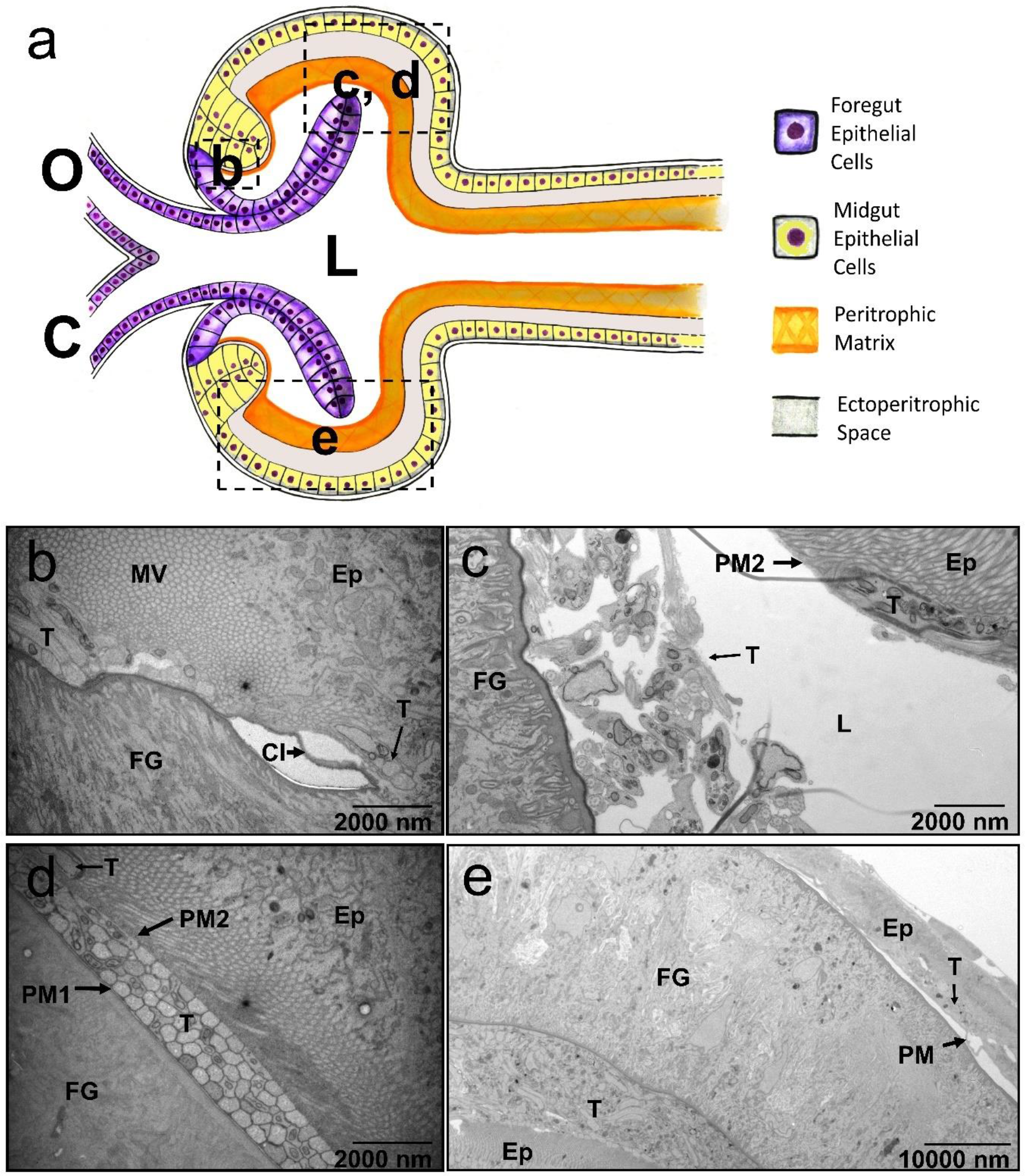
TEM images of an early (5 dpi) proventricular invasion by trypanosomes. **a**, Schematic depiction of the tsetse proventriculus as seen in the sagittal plane. The proventriculus is a transition area between the foregut (purple), comprised of the oesophagus (O) and crop (C) (lined with cuticular intima, CI), and midgut cells (yellow). The PM (orange) originates from a number of specialised epithelial cells, collectively termed annular pad, and is continuously secreted posteriorly. Dashed squares represent the approximate areas that the micrographs in figures 5b-e were taken from, with the letters inside corresponding to the lettered micrographs. L, lumen. **b**, Area of cell transition between the foregut (FG) and the proventricular/midgut epithelial cells (Ep). Trypanosomes (T) can already be seen near to the epithelial cells. CI, Cuticular Intima. MV, microvilli (8200 ×). **c**, Only PM2 is visible and trypanosomes appear to be filling the cavity between the foregut and epithelial cells (8200 ×). **d**, Tightly packed parasites can be seen already trapped between PM1 and PM2, and a single trypanosome (T) can also be seen already in the ES (8200 ×). **e**, The proventriculus is heavily infected, and parasites are located in the lumen, the ES and between PM layers (1700 ×). All midguts from the same proventriculus samples shown in this figure had trypanosome infections (not shown).

**Fig. 5.**
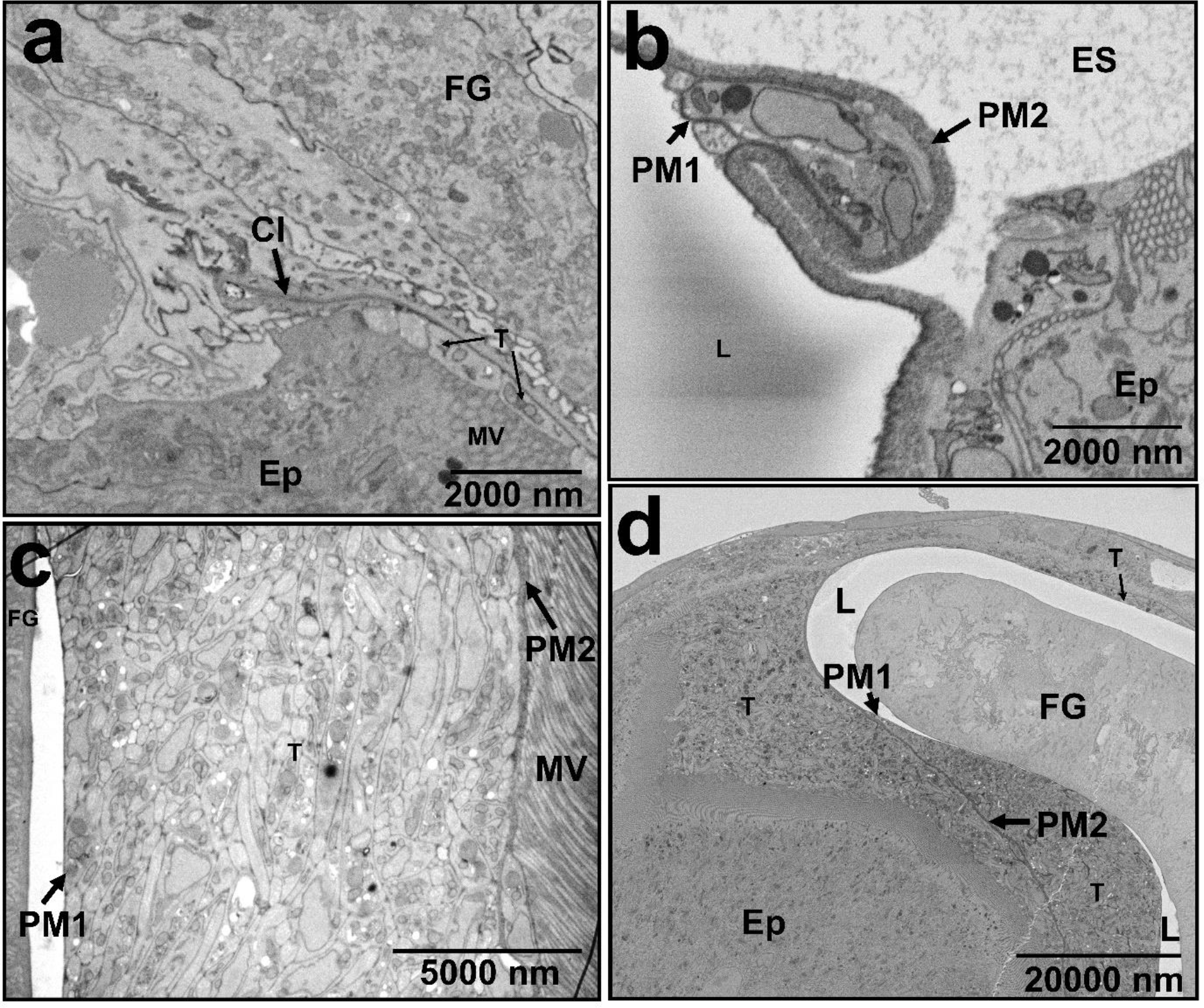
Proventricular trypanosomes contained within the PM and formation of trypanosome-filled cysts at 11 dpi. Micrographs are taken from an equivalent area of the proventriculus as shown in Fig. 5. **a**, Area of cell transition between the foregut (FG) and the epithelial cells (Ep). Trypanosomes (T) can already be seen near to the epithelial cells. CI, Cuticular Intima. MV, microvilli (8200 ×). **b**, Cysts of trypanosomes are formed in the proventriculus (8600 ×). **c**, Trypanosomes neatly contained in the ES with no visible parasites in the lumen (L). Parasites can also be seen between PM1 and PM2 (1700 ×). **d**, High numbers of parasites (T) can be seen trapped between PM1 and PM2 layers, and within the ES (8600 ×).

At 11 dpi, trypanosomes continued to be seen in the proventriculus (Fig. 5) (50% prevalence). However, whilst at 5 dpi parasites were located in the ES, the lumen and also between PM layers, by 11 dpi trypanosomes were neatly contained either within PM layers or inside the ES (Fig. 5c-d). In addition, cyst-like structures such as those in the anterior midgut could be observed (Fig. 5b) and with no evidence of a damaged PM1 layer. In summary, at 5 dpi, flies show two clear phenotypes: susceptible – those that have a high parasite load (including trypanosomes near to the cuticular intima of foregut cells) and refractory – those with no sign of parasite infection. In the former, trypanosomes widely distribute throughout the proventriculus, filling all available spaces. The high parasite numbers indicate trypanosomes are replicating during early infection. In contrast, by 11 dpi, parasites are absent from the proventriculus lumen and concentrated within PM layers. Overall, TEM analyses of infected proventriculi suggest trypanosomes are capable of penetrating the PM at its point of synthesis. Here PM2 is not fully formed and exists as a disorganised structure [21], so it is possible for trypanosomes to become passively engulfed by it rather than actively penetrating as previously suggested.

### CLSM confirms early proventricular colonisation

To further demonstrate an early proventricular invasion by trypanosomes, we used CLSM to localise live parasites (eGFP-expressing J10 BSFs, one of the strains used for TEM analysis) within tsetse tissues over a 5-day time course (Fig. 6). This parasite strain expresses eGFP only upon transformation into procyclics. Trypanosomes were detected within the proventriculus from 2 dpi (10% infection prevalence) onwards; however, at 3 dpi (Supplementary Fig. 4), heavy proventricular infections could be seen in 35% of the flies. Additionally, most of the flies at 3-5 dpi presented a high midgut infection particularly in the anterior midgut, either within the ES or the midgut lumen (Fig. 6a and 6d, and Supplementary Videos 8, 11 and 12).

**Fig. 6.**
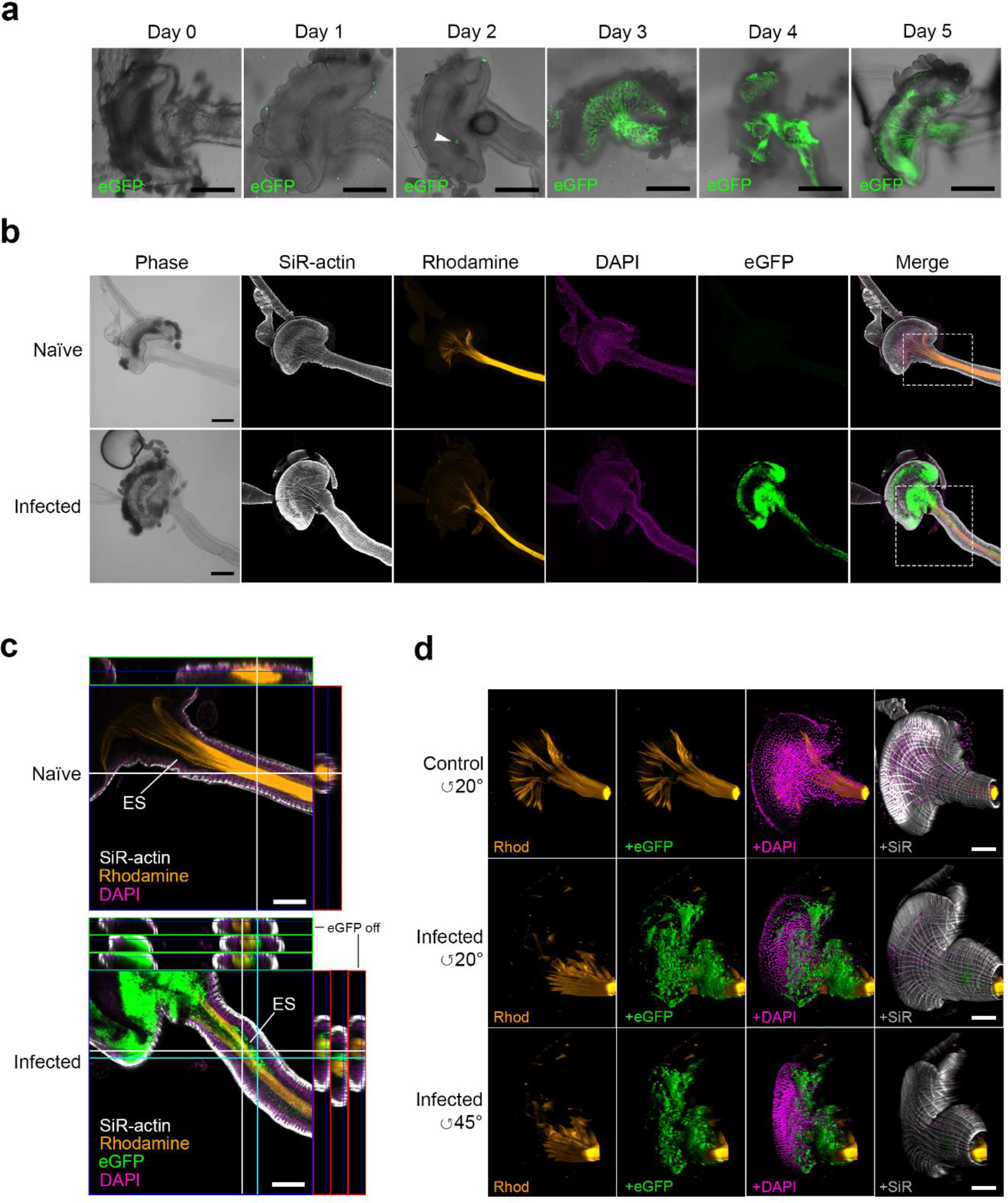
CLSM analysis of the early proventicular infection by bloodstream trypanosomes. **a**, Time course of infection up to 5 dpi with eGFP BSFs J10 strain. Figure shows representative proventriculi and anterior midguts from each dpi. “Day 0”, proventriculus from a fly dissected 1h after receiving an infected meal. White arrowhead at 2 dpi shows trypanosomes within the proventriculus (see also Supplementary video 8). Scale bar 200 μm under 10×. **b**, Example of an infected proventriculus and anterior midgut (3 dpi) showing the location of trypanosomes (green) in relation to the PM (orange). Top panel, naïve flies (Supplementary videos 9 and 10). Both naïve and trypanosome-infected flies received serum meals containing rhodamine-WGA four hours prior to dissection, which shows PM originating from proventriculus. SiR-actin labels the filamentous-actin (white) of all tsetse proventricular cells and DAPI (magenta) their nuclei. Bottom panel, a heavy trypanosome infection inside the proventriculus and anterior midgut (Supplementary videos 11 and 12). Insets were analysed at a higher magnification (6c). Scale bars 100 μm under 10X. **c**, **top**, Representative stack from a 3D-reconstructed proventriculus and anterior midgut at the region of interest from a naïve fly. Scale bar 100 μm under 10×. **c**, **bottom**, Representative stack from a 3D-reconstructed infected proventriculus and anterior midgut. Trypanosomes can be seen in the ES within either the proventriculus or the anterior midgut, whilst the orthogonal views show trypanosomes either within the lumen or the PM layers (white section), or within the ES (cyan section). Scale bar 100 μm under 10×. **d**, CLSM 3D reconstructions from multiple z-stacks of proventriculi and anterior midgut from a naïve (top) and infected fly at 5 dpi (middle and bottom). Scale bar 100 μm under 10×.

The same early proventricular infection phenotype was also seen in flies infected with BSFs from another *T. b. brucei* strain (AnTat 1.1, clone 90:13) (Supplementary Fig. 5a). However, and completely unexpected, when infections were carried out using *in vitro* cultured AnTat 1.1 BSFs (cBSF) proventricular trypanosomes could only be seen at 15 dpi or later (Supplementary Fig. 5a-b). Furthermore, the ability of cBSFs to colonise the proventriculus few days after infection was severely reduced as early as 9 days after adaption in culture when compared to BSFs (Supplementary Fig. 5a). Strikingly, whilst BSFs are able to establish normal salivary gland (SG) infections (21% infection prevalence), recently adapted cBSFs show lower prevalence (10%) and cBSFs completely fail (0%) to colonise the tsetse SGs (Supplementary Fig. 5a). Furthermore, when we retrospectively analysed infection data collected in our lab over a period of three years, it was confirmed that almost 30% of flies fed with AnTat BSFs developed SG infections (Supplementary Fig. 5c). In contrast, flies that received bloodmeals containing either procyclic or cBSFs produced ~10% and 0% SG infections, respectively, after 30 days. Thus, an early proventricular colonisation by bloodstream trypanosomes results in a higher infection prevalence of the tsetse salivary glands.

To understand the dynamics of trypanosome development in early proventricular infections, we isolated parasites from infected proventriculi and midguts at 5 and 15 dpi, and analysed length (Fig. 7), morphology, and kinetoplast position relative to the nucleus [4, 31, 33] (Supplementary Fig. 6). Proventricular trypanosomes at 5 dpi, although on average ~3μm shorter in length than those observed at 15 dpi, were significantly longer (~35μm average length) and morphologically different than midgut procyclics, at either time point. This implies procyclic forms can differentiate into mesocyclics early on within the proventriculus. Epimastigote forms developed at a slower rate and were only detected from 15 dpi.

**Fig. 7.**
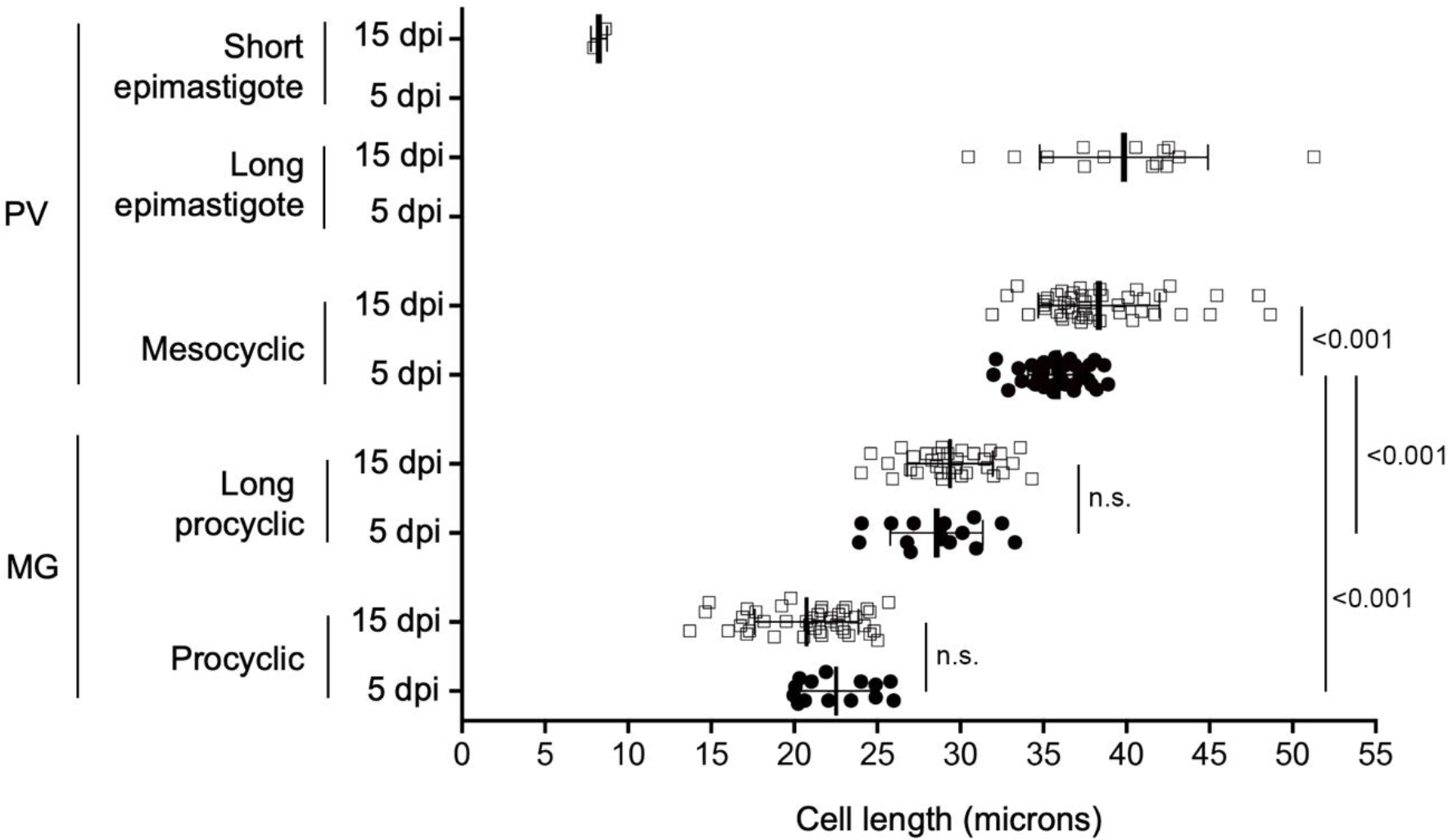
Analysis of trypanosome life stages at different infection times and tsetse tissues. Mean cell length (μm) of trypanosomes isolated from fly midguts (MG) and proventriculi (PV) at either 5 dpi (●) or 15 dpi (□) were DAPI stained and analysed by CLSM. Midgut trypanosomes include free swimming and encysted parasite forms. Error bars represent ± s.d. Vertical lines show statistical significance (one-sided *t-*test, assuming normal distribution) among life stage groups and the two time points (p-values indicated next to vertical lines).

We also compared the expression of procyclin, a surface glycosylphosphatidylinositol (GPI)-anchored glycoprotein marker, in proventriculus and midgut trypanosome populations using antibodies specific for each form (EP or GPEET; Fig. 8) [34]. We observed a similar pattern of procyclin expression in parasites isolated from both organs at 3, 5 and 7 dpi. Whilst EP procyclin was detected in 100% of cells at all time points, both forms of GPEET (unphosphorylated and phosphorylated) were primarily detected in proventricular and midgut forms at 3 dpi. Altogether, these results suggest that although proventricular trypanosomes may be developing at a faster rate than those in the midgut, the programme of procyclin expression mirrors that of proliferating midgut procyclics; i.e. GPEET is only expressed early on during the infection (regardless of the trypanosome stage and tissue infected) and EP becomes the dominant form from 5 dpi onwards [2, 3]. Interestingly, at 3 dpi, midgut trypanosomes showed a fully posterior kDNA compared to proventricular forms at the same time-point (Fig. 8c), which is more reminiscent of transforming ‘stumpies’ than fully developed procyclic cells.

**Fig. 8.**
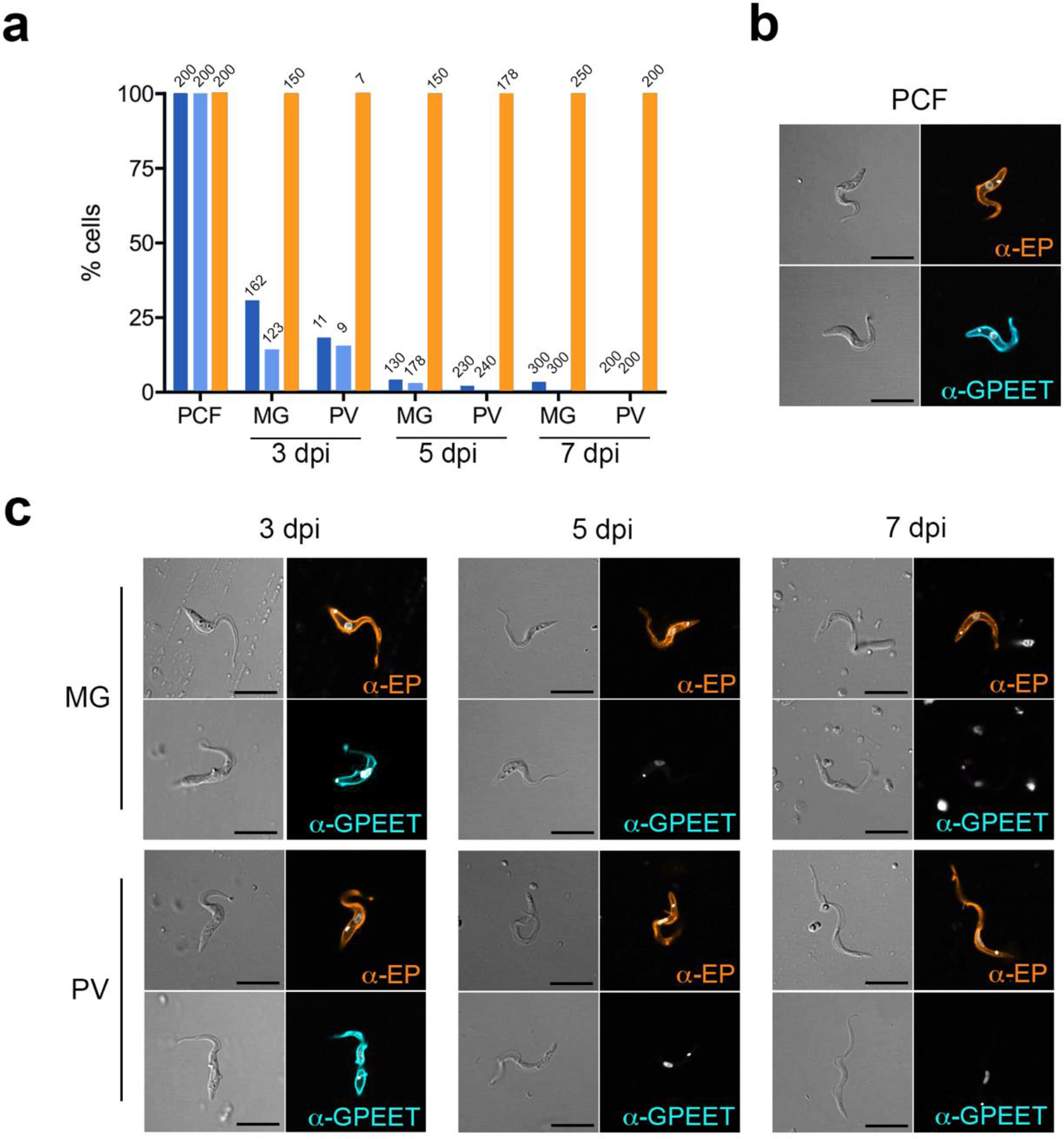
*T. brucei* procyclin expression during early infection in the tsetse. **a**, Profile of procyclin expression in trypanosomes during a time course infection experiment (*n*=1) as determined by immunostaining. Percentage of *T. brucei* cells from midgut (MG) and proventriculus (PV) at 3, 5 and 7 dpi, either recognised by antibodies against GPEET (phosphorylated (dark blue) or unphosphorylated (light blue)) or EP procyclins (orange). Numbers on bars represent individual trypanosomes analysed. **b**, Representative immunostaining images of *T. brucei* procyclic cultured forms (PCF) (antibody controls) shown in differential increased contrast (left) and a merged image of DAPI DNA counterstain (white) with either anti-EP (orange) or anti-GPEET (phosphorylated; blue) (right). **c**, >100 cells per tissue and time point were analysed for each antibody with the exception of cells from the PV at 3 dpi due to very few infections at this time (*n*=11, 9 and 7 for phosphorylated GPEET, unphosphorylated GPEET and EP, respectively). Imaged at 63×; Scale bars 10μm.

### Do serum factors influence early colonisation of the proventriculus?

We also investigated whether factors in trypanosome-infected serum promoted an early proventricular colonisation (Fig. 9). Teneral flies that received bloodmeals consisting of established AnTat 1.1 90:13 cBSFs, spiked with either naïve serum or serum from mice originally infected with BSFs AnTat 1.1 90:13, showed no proventricular colonisation compared to BSFs at 5 dpi (i.e. >30% infection prevalence; Fig. 9a). Infectivity was only evident at 10 dpi (Fig. 9b). Surprisingly, flies that were fed with bloodmeals consisting of AnTat 90:13 procyclic cultured forms (PCFs) spiked with infected serum showed a significant 10.3-fold increase in proventriculus infection prevalence (>87% average) at 5 dpi compared with the control serum group. This suggests that serum factors from trypanosome-infected blood may facilitate early proventricular infections only once transformation from BSFs into PCFs has occurred within the fly gut. However, these results also indicate that intrinsic cell factors are important to establish an early proventricular infection as this phenotype was lost during long-term culture and could not be rescued in the presence of trypanosome-infected serum (Fig. 9a).

**Fig. 9.**
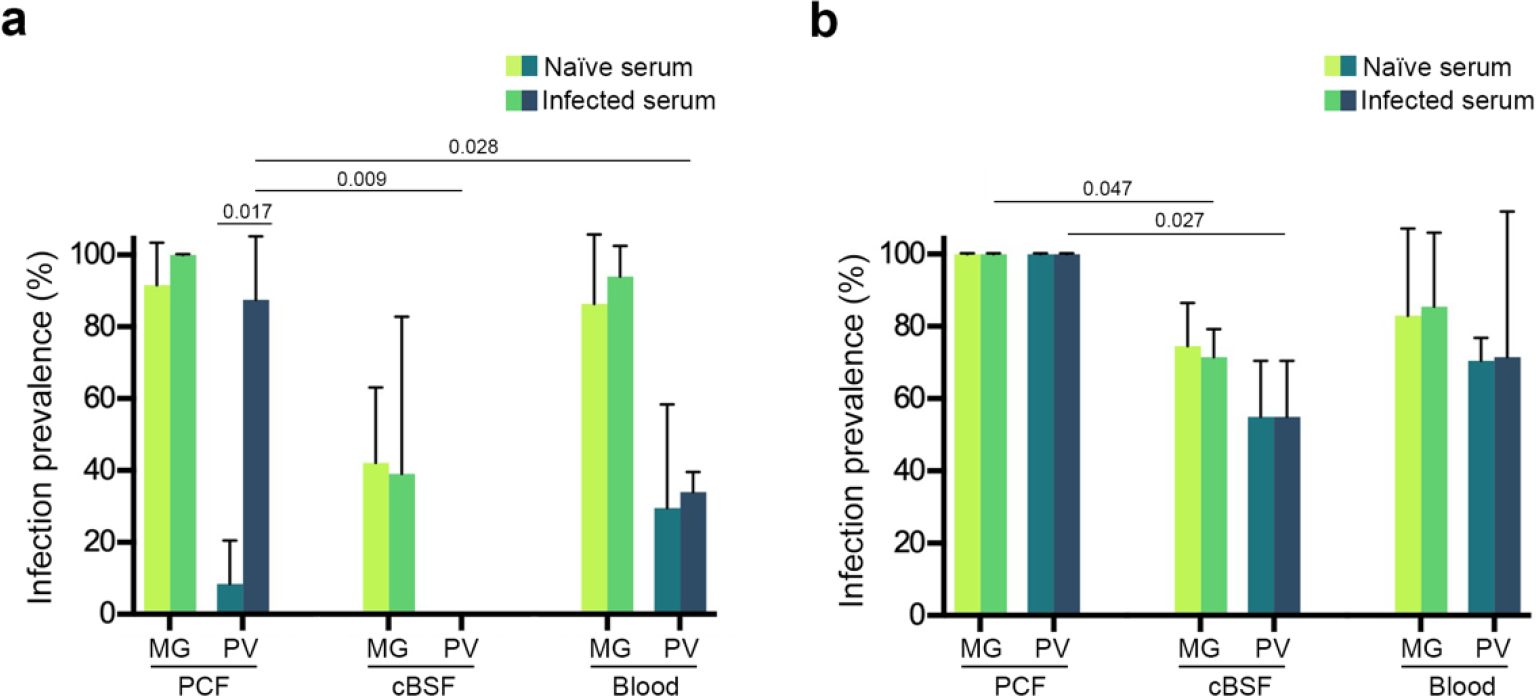
*T. brucei* life stages display different infection phenotypes in the fly when in the presence of serum from trypanosome-infected animals. Mean trypanosome infection prevalence (percentage) in midguts and proventriculi of tsetse infected with bloodmeals consisting of serum harvested from either trypanosome-infected blood or naïve and then equally combined with washed parasites (either cultured procyclics forms (PCFs), cultured bloodstream forms (cBSFs) or bloodstream forms from infected mice (BSFs)) and horse blood, were given to teneral flies and then infection prevalence was scored after 5 (**a**) or 10 (**b**) dpi. Error bars show ± s.d. Horizontal lines represent statistical significance from two biological replicates (*n*=2) using a one-sided *t-*test assuming normal distribution (p-values indicated on the lines).

One possible serum factor that could promote establishment of an early proventricular infection are released variable surface glycoproteins (VSGs) [35–39]. However, when flies were infected with Antat 1.1 PCFs in bloodmeals containing several concentrations of soluble (i.e. GPI-cleaved) VSG variant MITat1.4, we saw no significant difference in either proventricular or midgut infectivity (data not shown). It is worth mentioning that upon ingestion of a trypanosome-infected bloodmeal, released VSG molecules –presumably from transforming parasites– lead to a transcriptional down-regulation of PM associated genes expressed by proventricular epithelial cells, including several peritrophins [29]. The authors conclude that this interference facilitates the crossing of the PM by procyclic trypanosomes in the anterior midgut during early infection. However, based on the data herein presented, we suggest that the VSG-induced down-regulation of proventricular genes may instead facilitate parasite crossing of the PM in the proventriculus rather than in the anterior midgut in which the PM is present as a fully assembled, multi-layered tissue. Furthermore, whilst at a transcriptional level this may be true, a comparison of the PM width in the anterior midgut between naïve and infected flies (at either 5 or 11 dpi) showed no significant difference in architecture or thickness as evaluated using TEM. On average, the tsetse PM is ~300nm in all conditions (Supplementary Fig. 7).

Our results support a new infection model where recently transformed *T. brucei* procyclics reach the ectoperitrophic space after crossing the peritrophic matrix located within the proventriculus [18, 19], and not in the anterior midgut as previously suggested [22, 24, 25]. In this scenario, procyclic trypanosomes can either first establish a proventricular infection and then gradually invade the ES after 3 dpi or, alternatively, directly establish an ES infection in the anterior midgut and then migrate to the proventriculus as the infection progresses (usually after one week depending on the parasite strain). The precise location of PM penetration by trypanosomes within the proventriculus remains unknown; however, we hypothesise that it may occur in the region where specialised epithelial cells (annular pad) secrete the different PM layers, as suggested by Fairbairn in 1958 [20] and later by Moloo in 1970 [21]. If so, this implies a race against time for trypanosomes as they must escape the confines of the PM to gain entry into the ES before it matures to a point where they become trapped in cysts that move along the midgut (due to continuous PM secretion) and then become potentially eliminated in the hindgut (posterior end; Fig. 10). Indeed, TEM analysis of the hindgut from infected flies showed parasites with abnormal morphology (i.e. containing many intracellular vesicles and multiple flagella) and a damaged PM (Supplementary Fig. 8), suggesting a possible degradation as parasite cysts transit towards this region. The remarkable PM expansion observed in some of the cysts at 11 dpi, where the PM layers remain intact despite containing several tightly packed trypanosomes, is indicative of a highly flexible PM rich in β-chitin fibres cross-linked to *O*-glycosylated peritrophins [14, 40]. Whether parasite encapsulation within the PM is a novel tsetse defence mechanism to control trypanosome infection intensity remains to be elucidated, but one should note this process does not lead to self-cure. In fact, in flies at least 40 dpi, similar cysts have been reported within the proventriculus in which the integrity of PM1 seems to be compromised, although the phenotypic differences could be accounted for by fly age and duration of infection [41]. In summary, the model of trypanosome PM crossing in the anterior midgut, which for several decades has been mainly supported by TEM visualisation of parasite cysts in the same region [22], can no longer be accepted as the sole route of ES invasion based on our collective microscopy evidence.

**Fig. 10.**
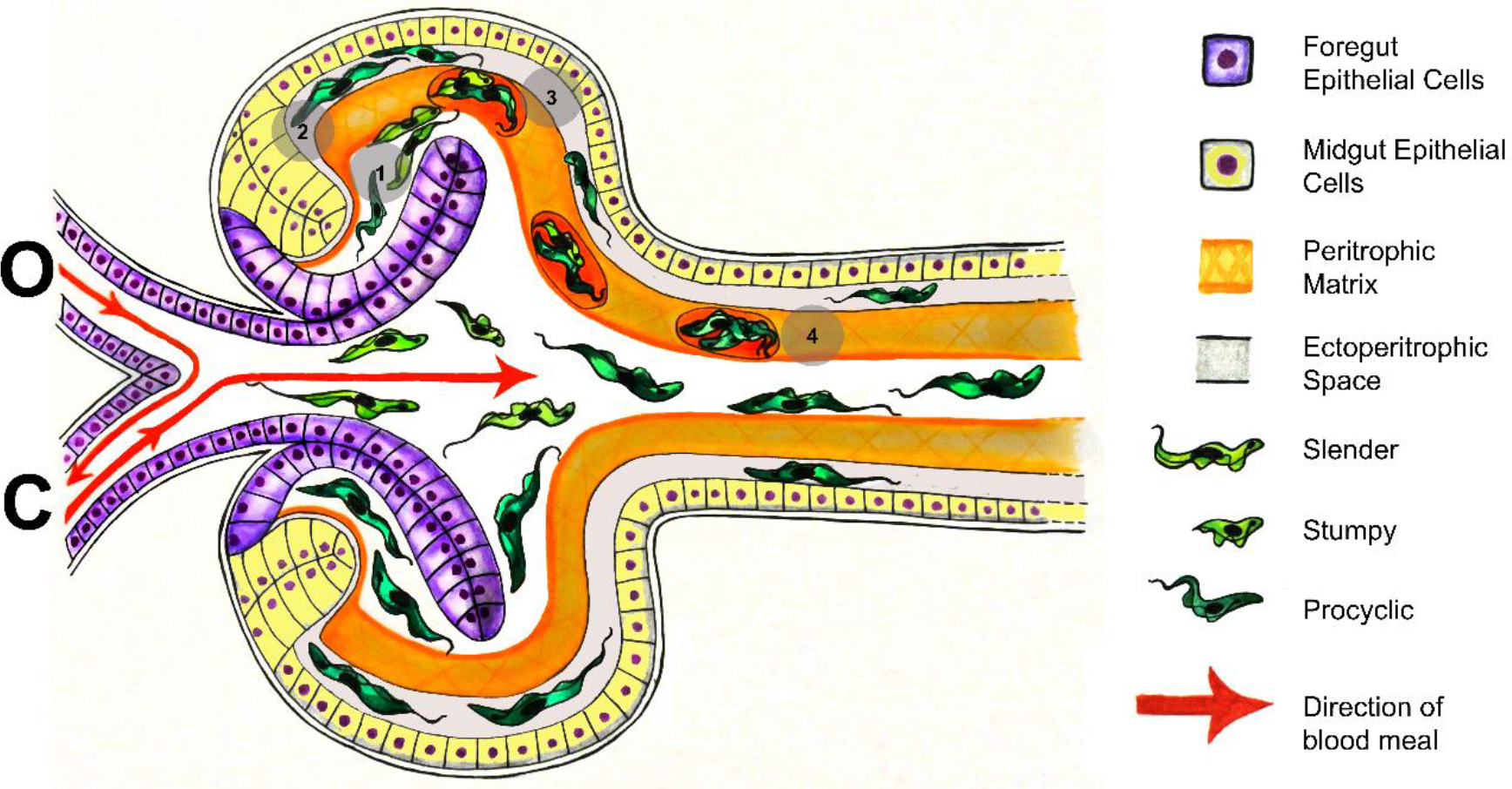
Schematic (artistic) representation depicting the entry trypanosomes into the ectoperitrophic space via the proventriculus. Specialised epithelial cells in the proventriculus annular pad are responsible for PM (orange) assembly and secretion. Ingested trypanosomes (green) either remain in the proventriculus lumen (1) successfully migrate to the ES through a more fluid PM in the proventriculus (2) before it maturates into a rigid structure as seen in the midgut, or become trapped between PM layers (cysts) (3). Those that have become trapped between PM layers are carried through to the midgut as the PM continues to be secreted (4). “O” and C” represents direction of the blood flow from either the oesophagus or crop, respectively.

Why procyclic trypanosomes ‘hide’ within the proventriculus and/or midgut ES to establish an infection is unknown. As previously suggested, the most likely explanation is that the tsetse ES offers a safer environment to proliferative procyclic trypanosomes against the action of harmful blood factors, including reactive oxygen species [11, 42] and vertebrate complement [43]. In addition, considering that the tsetse PM is continually secreted, attachment to it from the midgut lumen would result in the eventual excretion of trypanosomes. This contrasts to mechanisms used by *Leishmania* and *Plasmodium* parasites within the gut of the sand flies and mosquitoes, respectively, as these parasites secrete chitinases to degrade the type I PM of these insects in order to migrate [26, 27, 44]. Trypanosomes do not secrete chitinases, however, exochitinase activity from the tsetse symbiont *Sodalis glossinidus* [45] or bloodmeal chitinases [46] may facilitate trypanosomes penetration into the proventriculus. Thus, invasion of the ES by penetrating an immature PM in the proventriculus may be an adaptive strategy to compensate for the inability of all *T. brucei* sub-species (and possibly also for *T. congolense*) to attach to and degrade a mature tsetse PM. In fact, a type II PM is a more complex and organised structure compared to type I PMs, and blood feeding insects secreting type I PMs are usually more permissive disease vectors [7].

It is not clear what the impact of an early proventricular colonisation has on trypanosome development or transmission. Our results indicate that establishment of an early proventricular infection may increase parasite transmissibility as the proportion of infected SGs is much higher compared to strains (or parasite stages) that first colonise the midgut. The tsetse proventriculus, besides regulating the blood flow coming from the oesophagus and crop, and also being the place of PM synthesis, is an immunoregulator organ that responds to a trypanosome infection by increasing the levels of nitric oxide and radical oxygen species, and parasite-specific antimicrobial peptides [11, 47]. Collectively, these molecules appear to be key in conferring to tsetse refractoriness to a trypanosome infection. Thus, it is possible that during an early colonisation of the proventriculus, procyclic trypanosomes in combination with serum factors present in infected blood down-regulate the release of immunoregulator molecules, which in turn will facilitate establishment of a parasite infection and a faster development (i.e. formation of epimastigotes) within this organ. This is in contrast to a later proventricular colonisation phenotype, which normally occurs after 10 dpi and correlates with a lower transmission index.

There is increasing evidence that procyclic trypanosomes undergo social motility (SoMo) *in vitro* [48–50]. This phenomenon appears to occur only in early procyclic cells, which are characterised by expressing GPEET procyclin on the surface [51]. Furthermore, although SoMo is yet to be observed within the fly’s midgut, it may play a role in the migration of midgut procyclics to the proventriculus [52, 53]. We did not investigate whether trypanosome SoMo occurs *in insecta*, but our data suggest that this phenomenon could happen in developing (early) procyclics in the proventriculus, as supported by the expression of GPEET procyclins in proventricular-associated parasites (Fig. 8). Alternatively, there is strong evidence for trypanosome CoMo within infected tsetse tissues [31], although this may not be operative during an early proventricular infection. However, both phenomena (SoMo *vs*. CoMo) are not necessarily mutually exclusive as they could operate in parallel or at different stages of trypanosome development in tsetse.

In conclusion, we have developed new microscopy methodologies that allowed us to revisit the route by which trypanosomes migrate through the tsetse gut. We provide evidence that *T. brucei* procyclics reach the tsetse ES when they encounter the immature PM secretions at its point of production in the proventriculus. Furthermore, trypanosomes observed within PM cysts in the anterior midgut are likely formed in the proventriculus during PM assembly and are not indicative of PM penetration in this region. Moreover, unknown factors present in infected blood (of mammalian and/or parasite origin) may promote early proventricular invasion, which in turn leads to higher salivary gland infection rates and potentially increasing parasite transmission.

## Materials and Methods

### Tsetse

Male flies were reared in an established colony (*Glossina morsitans morsitans* (Westwood)) at the Liverpool School of Tropical Medicine and maintained on sterile, defibrinated horse blood (TCS Biosciences) at an ambient temperature of 27°C ± 2°C and a relative humidity of 65-75%. Experimental flies were collected at <24 hours post-eclosion (p.e.) and offered a bloodmeal every 2 days before being starved for 72 hours in preparation for dissection at variable timepoints (5, 8 or 11 dpi) control blood meal. Flies used for CLSM were an exception to this feeding regime (see below).

### Trypanosome strains

Three different strains of *Trypanosoma* (Trypanozoon) *brucei brucei* were used in this study. TSW-196 BSFs [54] (from murine stabilates) were used for TEM experiments. J10 (MCRO/ZM/73/J10) green fluorescent protein (eGFP) expressing BSFs [55] were used for TEM, CLSM and both BSFs and procyclic forms (PCFs) used in the procyclin expression experiments. BSFs of AnTat 1.1 90:13 engineered with an mNeonGreen expressing construct was used for CLSM (see below), whilst cultured BSFs (cBSFs) and PCFs of the same strain was used for CLSM and infected serum experiments. For infections, flies <24 hours p.e. were fed either a blood or serum meal containing one of the strains described above; unfed flies were removed and conditions prior to sacrifice are the same as described for control flies.

### Trypanosome growth and transformation

Cultured BSFs were grown in HMI-9 supplemented with 10% foetal bovine serum (FBS) at 37°C with 5% CO_2_ whereas PCFs were grown in SDM-79 with 10% FBS at 27°C and 5% CO_2_. Cultured BSFs were transformed to PCFs using 6mM cis-aconitate in DTM (Differentiation medium [56]) supplemented with 20% FBS at 27°C and 5% CO_2_ for 24 hours. To generate the trypanosome mNeonGreen clone, 4×10^7^ AnTat 1.1 90:13 cultured BSF cells in exponential growth phase were transfected with 10μg of a modified pALC14 plasmid ([57]) for ectopic expression of the tetracycline-inducible mNeonGreen protein under a *GPEET* procyclin promoter, using an Amaxa 4D nucleofector (program FI-115). Clonal cell lines were selected by limiting dilution in SMD-79 10% FBS, containing 1μg/mL puromycin.

### Transmission Electron Microscopy

Tsetse midguts or proventriculi were dissected in ice-cold fixative (0.2M cacodylate, 4% paraformaldehyde (PFA), 2.5% glutaraldehyde (GA), 3% sucrose, pH 7.4), transferred to fresh fixative, and incubated on ice for an hour. Tissues were then washed twice in ice-cold 0.1M cacodylate buffer containing 3% sucrose (pH 7.4) for 2 minutes and left in 1% osmium tetroxide for an hour at room temperature. After washing with copious amounts of ice-cold 0.1M cacodylate buffer, followed by washes with distilled water, tissues were placed in 0.5% uranyl acetate in 30% ethanol before going through a series of 10 minute ethanol washes in increasing concentration (30-80%) and left for 30 minutes in 100% ethanol. Graded hard embedding resin 182 (TAAB) was mixed in a 1:1 ratio with 100% ethanol and left on tissues overnight, then replaced with fresh 100% resin for 30 minutes and placed in an oven at 60°C for 48 hours to cure. Ultrathin orthogonal serial sections (70-74 nm) were cut through regions of interest and collected on freshly prepared Pioloform®-coated 200 (for midguts and proventriculi) or 100 (for proventriculi) mesh nickel grids, before post-staining in uranyl acetate (5% w/v in 30% ethanol) and 50% lead citrate. Sections were viewed at 100 KV in a FEI Tecnai G2 Spirit and all micrographs were taken using either an Olympus Megaview3 or a Gatan Orios camera with AnalySIS or Gatan GMS2 software respectively.

### SBF-SEM and 3D reconstructions

Tissues were prepared and stained for SBF-SEM and 3D reconstruction using a modified method based on the protocol of Deerinck et al 2010 [58]. Briefly, tsetse midguts were dissected in ice-cold fixative (0.1M cacodylate, 2% paraformaldehyde (PFA), 2% glutaraldehyde, 3% sucrose, 2mM calcium chloride pH 7.4) or modified fixative (0.1M cacodylate, 2% PFA, 2% GA, 3% sucrose, 0.1% tannic acid pH 7.4) followed by washes in 0.1M cacodylate buffer pH 7.4 with 2mM calcium chloride prior to staining with reduced osmium tetroxide (2%) containing 1.5% potassium ferrocyanide in 0.1M cacodylate buffer. Midguts were washed in distilled water and incubated in 1% thiocarbohydrazide (TCH) for 30 minutes before further washes in distilled water. A second osmium (2%) staining was carried out at room temperature for 40 minutes, followed by washing in distilled water before incubation in 1% aqueous uranyl acetate overnight at 4°C. A final wash in distilled water was carried out and samples were stained in warmed lead aspartate for 30 minutes before dehydration in graded ethanol 30-90% followed by 100% ethanol. For samples dissected in modified fixative an additional step was added and samples were placed in 100% propylene oxide following the series of ethanol washes. Samples were placed in hard resin 812 (TAAB) at a 1:1 ratio with 100% propylene oxide and left overnight before infiltration with increasing ratios of resin:propylene oxide until 100% resin and left to cure for 48 hours. Samples were prepared for SBF-SEM by mounting a small square of embedded sample onto a cryo pin with conductive epoxy. Excess resin was trimmed away with an ultra-microtome and the sample coated with 10nm AuPd using a Q150T sputter coater (Quorum Technologies). Samples were imaged with a FEI Quanta 250 FEG modified with a Gatan 3View running GMS2 software. All samples were imaged in Low vacuum mode with a chamber pressure of 50 Pa. For the midgut imaging conditions were 2.2 KV, dwell time of 12 μs per pixel, magnification 26 K, giving a resolution of 3.3 nm in X and Y 100nm in Z over 474 slices of which the first 200 were taken for reconstruction. For proventriculus imaging conditions were 2 KV, dwell time 12 μs per pixel, magnification 8.7 K and Z was reduced to 40nm and 3 regions of interest (ROI) were scanned, all of which were 458 with a resolution of 18.7 nm in X and Y. The reconstruction was carried out on the first 400 slices of ROI 2. GMS2 was used for alignments and conversions to TIFFs and the reconstructions were carried out using Bitplane Imaris version 8.1.

### Confocal Laser Scanning Microscopy

#### Whole tissues

*G. m. morsitans* teneral flies (0-24 hours old) were infected with FBS containing 10% rat blood spiked with either cBSF or BSF AnTat or J10 eGFP BSF trypanosomes (final density of 2 × 10^6^ cells) and 10μg/mL wheat germ agglutinin (WGA)-rhodamine. A naïve group (uninfected) was fed with serum meal only. Flies were fed every day with FBS 10μg/mL WGA-rhodamine and dissected in PBS to score trypanosome infection in midgut and proventriculus. The proventriculi were fixed in fresh 1% PFA on ice for 1 hour, stained with SiR-actin (1:1000 dilution in PBS, Cytoskeleton Inc.) for 4 hours, incubated with 300ng/mL DAPI for 10 minutes and mounted in 1% low melting agarose with SlowFade Diamond antifade (ThermoFisher). Samples were imaged using a Zeiss LSM800 confocal laser scanning microscope.

#### Isolated PM

Flies infected with J10 eGFP BSF trypanosomes, were dissected at 5, 9 or 11 dpi and their peritrophic matrix was dissected out in ice-cold fresh 1% PFA and transferred to poly-lysine slides for 1 hour. Samples were incubated with 10μg/mL WGA-rhodamine and 300ng/mL DAPI for 15 minutes, washed and mounted in SlowFade Diamond antifade (ThermoFisher). Samples were imaged using a Zeiss LSM800 confocal laser scanning microscope.

#### Procyclin immunostaining

Teneral flies were infected with BSFs J10 eGFP strain and after 3, 5 or 7 dpi both proventriculi and midguts were dissected on a glass slide in fresh PBS and each tissue manually ruptured. Released parasites were harvested and pooled for each tissue and timepoint. Cells were gently pelleted and fixed in 4% PFA for 30 minutes, before washing in PBS and added to poly-lysine slides. Cells were left to adhere for 30 minutes at room temperature in a humid chamber before an hour block in 20% foetal bovine serum in PBS. The following anti-procyclin antibodies were then added for 1 hour in blocking solution; mAb 9G4 (mouse anti-GPEET unphosphorylated form, Biorad) 1:200 dilution, mAb 5H3 from hybridoma supernatant (mouse anti-GPEET phosphorylated form, Professor Terry Pearson) 1:10 dilution and mAb Clone TBRP1/247 (mouse anti-EP, Cedarlane) 1:800 dilution. The secondary antibody, anti-mouse IgG conjugated to Alexa Fluor 555 (ThermoFisher) was used at a 1:1000 dilution in blocking solution for an hour followed by 300 ng/mL of DAPI (ThermoFidsher) for 10 minutes. Samples were mounted in SlowFade Diamond antifade (ThermoFisher) and imaged using a Zeiss LSM800 confocal laser scanning microscope. Cultured AnTat 1.1 90:13 PCFs were used as antibody positive controls.

#### PM thickness measurements

100 different images from different flies and separate experiments for each time point and group was used: 5 dpi, 5-day naïve, 11 dpi and 11-day naïve. Each image from each time point/group was overlaid by a 10×10 square grid and a random number generator (numbers between 1-10 only) used to determine X and Y squares in which to take measurements. Measurements were made using ImageJ [59].

## Acknowledgements

We thank Professor Terry Pearson (University of Victoria, Canada) for providing us with anti-GPEET mouse hybridomas, Dr Lúcia Güther and Professor Mike Ferguson (University of Dundee) for the generous supply of *T. brucei* sVSG MiTat1.4 variant, Professor Sue Vaughan (Oxford Brookes University) for making available essential TEM protocols, and Prof Wendy Gibson for kindly supplying the *T. brucei* J10 strain. We thank Dr Lee Haines for critical reading of the manuscript, Dr Laura Jeacock for artwork and members of the Acosta Serrano group for constructive discussions. This work was supported by Wellcome Trust project grant 093691/Z/10/Z (awarded to AAS), Multi-User Equipment Grant (for confocal images 104936/Z/14/Z), GlycoPar-EU FP7 Marie Curie Initial Training Network (GA. 608295) (awarded to ACS and AAS), MRC Concept in Confidence Award MC_PC_17167 (awarded to AAS) and a PhD studentship from LSTM (awarded to CR).

## Competing interests

The authors declare no conflict of interest.

## Author contributions

CR, NAD, ACS, MJL, and AAS conceived and designed experiments. CR, NAD, AJB, BM, CS, MJL and IAP conducted and analysed EM work. ACS, CR, NAD and MM obtained confocal data. CR and AAS wrote the paper with input from all authors.

