## Supplementary data for "*Trypanosoma brucei* colonises the tsetse gut via an immature peritrophic matrix in the proventriculus"

[1]Department of Vector Biology and [2]Department of Parasitology, Liverpool School of Tropical Medicine, Liverpool, UK. [3]EM Unit, Department of Cellular and Molecular Physiology, Institute of Translational Medicine, University of Liverpool, Liverpool, UK. [4]Centre for Cell Imaging, Institute of Integrative Biology, University of Liverpool, Liverpool, UK. [5]Physiological Laboratory, Institute of Translational Medicine, University of Liverpool, Liverpool, UK.

‡These authors contributed equally.

†Present address: Department of Clinical Sciences, Liverpool School of Tropical Medicine, England, UK.

Contact during submission:

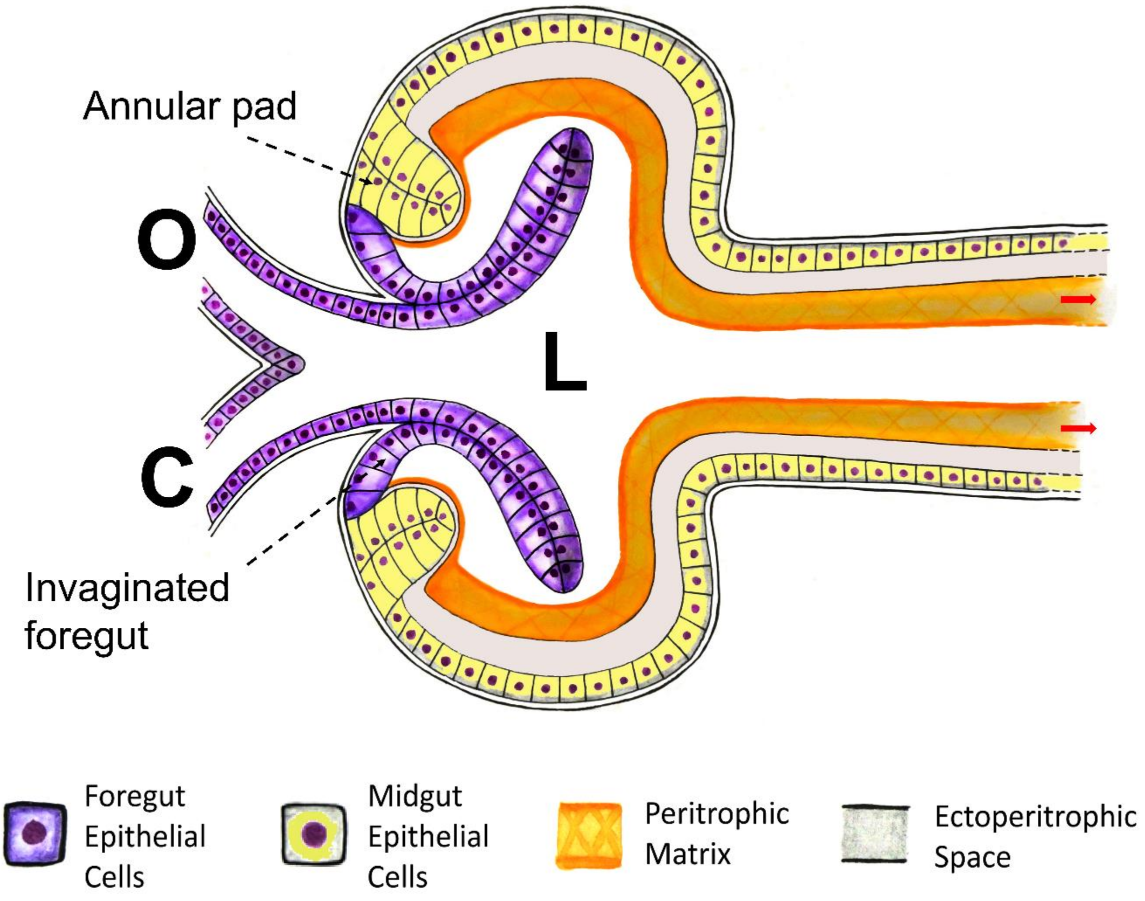

**Supplementary Fig. 1| Structure of the tsetse proventriculus.** Schematic depicting the proventriculus as seen in the sagittal plane. The tall columnar cells of the midgut (annular pad) curves around the extended, invaginated foregut (O, oesophagus C, crop) forming a press-like fold. The annular pad is responsible for PM secretion. The PM (orange) retains the food bolos inside the lumen (L) and red arrows indicate direction of formation as it matures and extends into the gut.

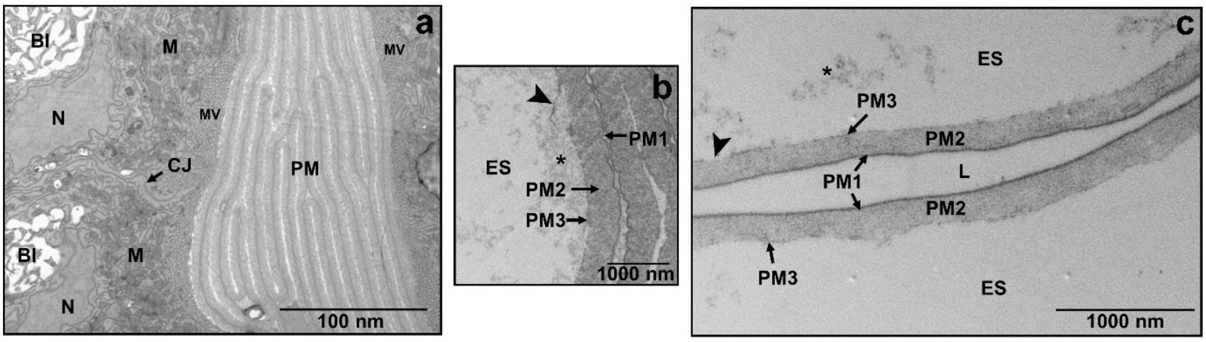

**Supplementary Fig. 2| Transmission electron micrographs showing naïve/refractory midguts and PM. a**, Shows the highly convoluted PM within the context of the gut in an unfed/starved fly. Taken from a fly at 11 days post-eclosion (naïve). BI; basal infoldings, M; mitochondria, MV; microvilli, N; nucleus, CJ; cell junction. **b**, A micrograph showing the 3 layers of the PM and secretions presumably coming from the epithelial cells (*) are being laid down onto PM2 (arrowhead). Taken from a fly at 11 dpi (refractory). **c**, A micrograph showing all 3 PM layers and few secretions (*) that stick onto the second layer. Along the length of the PM, PM3 is observed to be patchy (arrowhead). Taken from a 5 day old naïve fly.

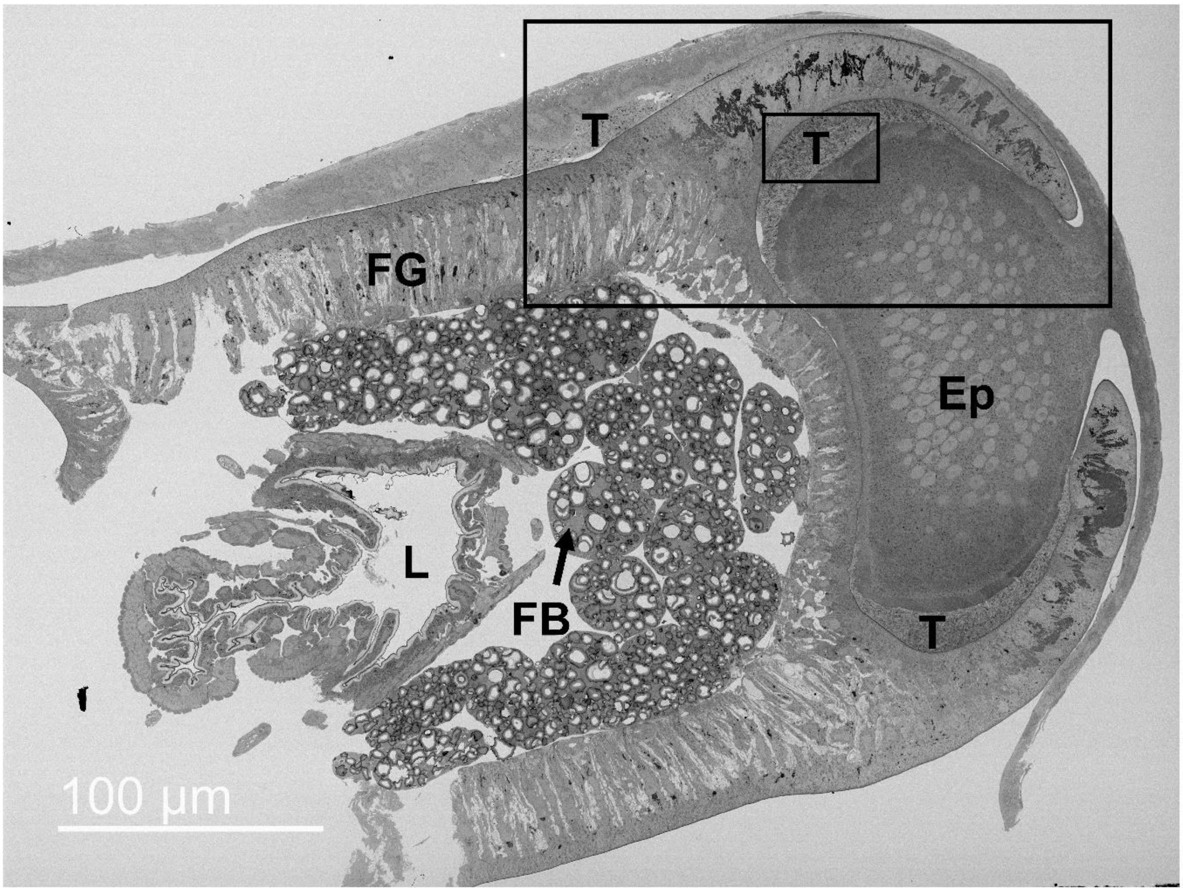

**Supplementary Fig. 3| An SEM micrograph showing the overview of an infected proventriculus at 5 dpi in a transverse plane.** Trypanosomes (T) can be seen occupying the spaces between the foregut and the epithelial cells of the proventriculus. The black boxes indicate the regions of interest (ROI) that were subject to SB-SEM (Outer box, ROI 1, Inner box, ROI 2). FB; Fat bodies, FG; Foregut, MG; Midgut, Ep; Epithelial cells

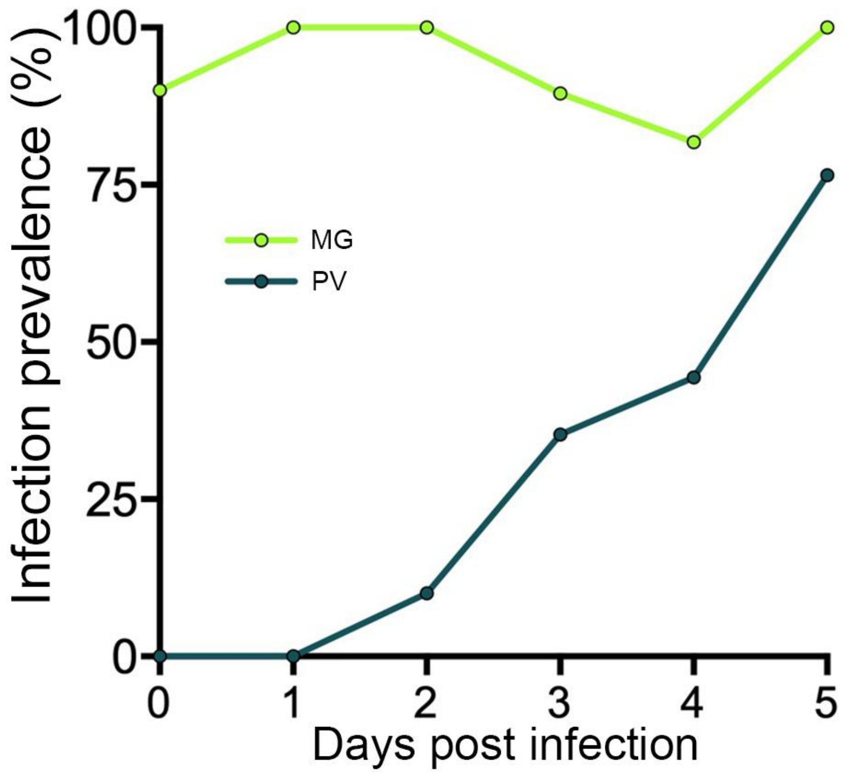

**Supplementary Fig. 4| Proportion of flies with a midgut or PV infection over 5 days.** Flies that had received a bloodmeal containing J10 BSFs (from murine stabilate) were dissected each day and midgut and proventriculus (PV) scored before preparation for LSCM.

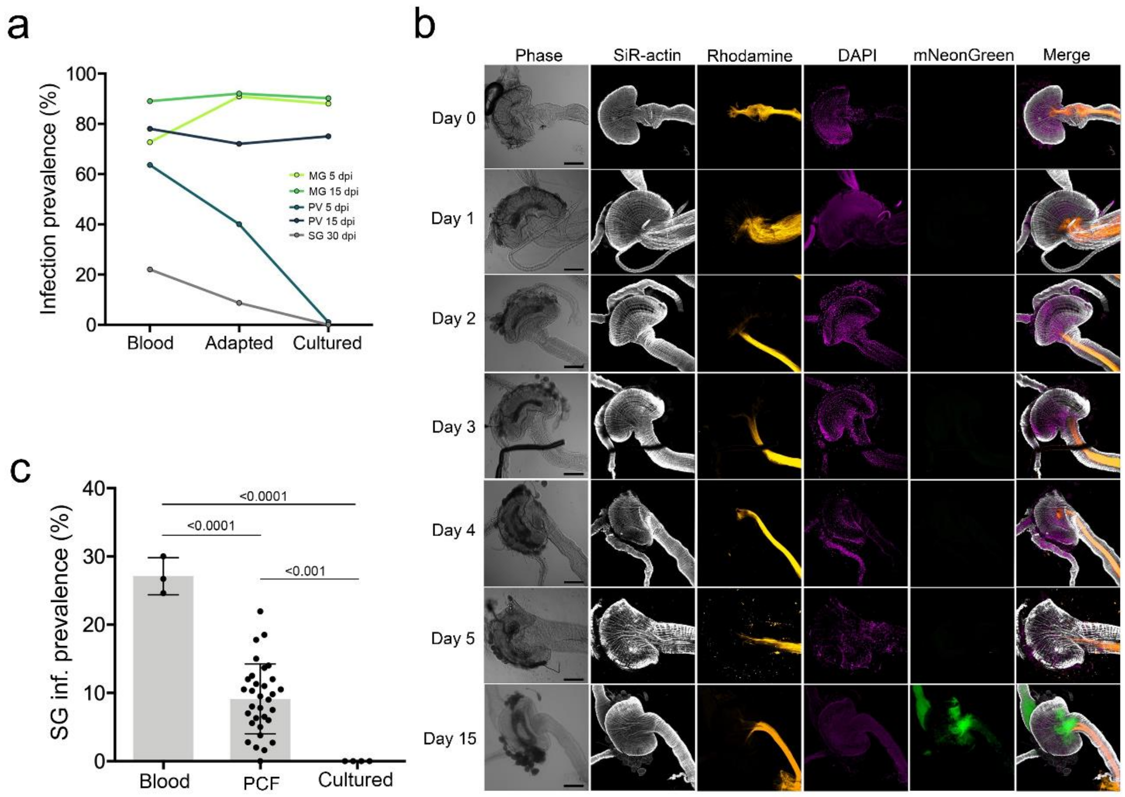

**Supplementary Fig. 5| Cultured bloodstream form (cBSF) trypanosomes gradually lose ability to invade the proventriculus early in infection leading to loss of salivary gland infections. a,** Flies were given an infected meal with either BSFs (blood), cBSFs that had been adapted to culture from the same BSF stabilate for just 9 days (adapted) and cBSFs that had been in culture for several months. Flies were dissected at 5 and 15 dpi and infection prevalence was scored for both the proventriculus (PV) and the midgut MG, or 30 dpi to check salivary gland (SG) infection. **b**, Flies were given an infected meal with cBSF *T. brucei* (Strain; AnTat 1.1 90:13 with a neon green construct) and dissected every day until trypanosomes were observed in the proventriculus. In contrast to BSFs from a murine stabilate, the cultured forms do not invade the proventriculus until at least 15dpi. The distribution is also different as very few trypanosomes can be seen in the ectoperitrophic space in the anterior midgut. Scale bar 100 µm, taken under 10X objective. **c**, Retrospective analyses of salivary gland (SG) infections using bloodstream forms (BSFs), procyclic forms (PCFs) or cultured BSFs (cBSFs). Each point represents independent experiments (with number of flies per experiment >100). All life stages used for all experiments were AnTat 1.1 90:13. Error bars represent ±s.d.; horizontal bars show statistical significance from a one-sided *t*-test assuming normal distribution; p-values are shown on the bars.

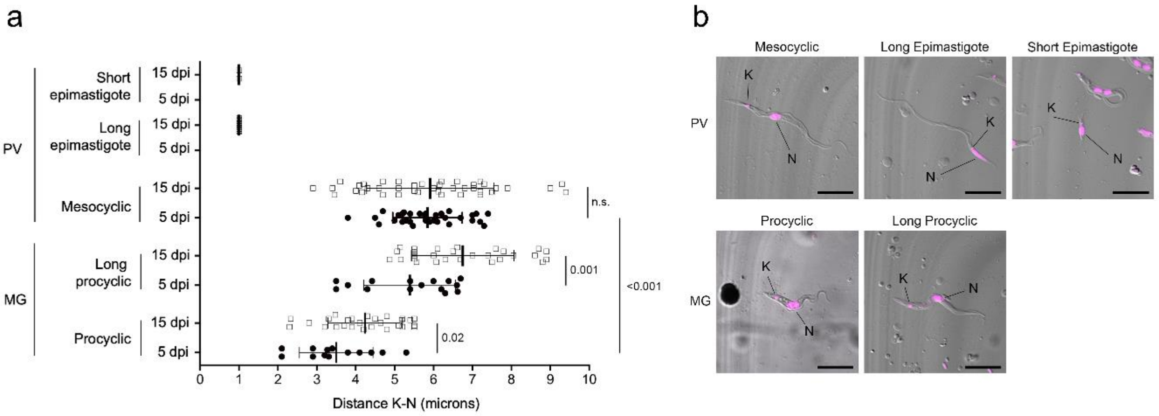

**Supplementary Fig. 6| Analysis of trypanosomes harvested from the proventriculus and midgut of flies 5 and 15 dpi. a**, Measurements of the distance between parasite nucleus and kDNA at different timepoints and tissues. **b**, Representative trypanosomes from each infected tissue. Trypanosomes shown in DIC with DNA (K, kinetoplast, N, nucleus) stained by DAPI (magenta). MG, midgut. PV, proventriculus. Scale bar 10µm. Error bars represent ± s.d. Vertical lines show statistical significance (one-sided *t*-test, assuming normal distribution) while comparing same parasite life stages among the two time points. p-values indicated next to vertical lines.

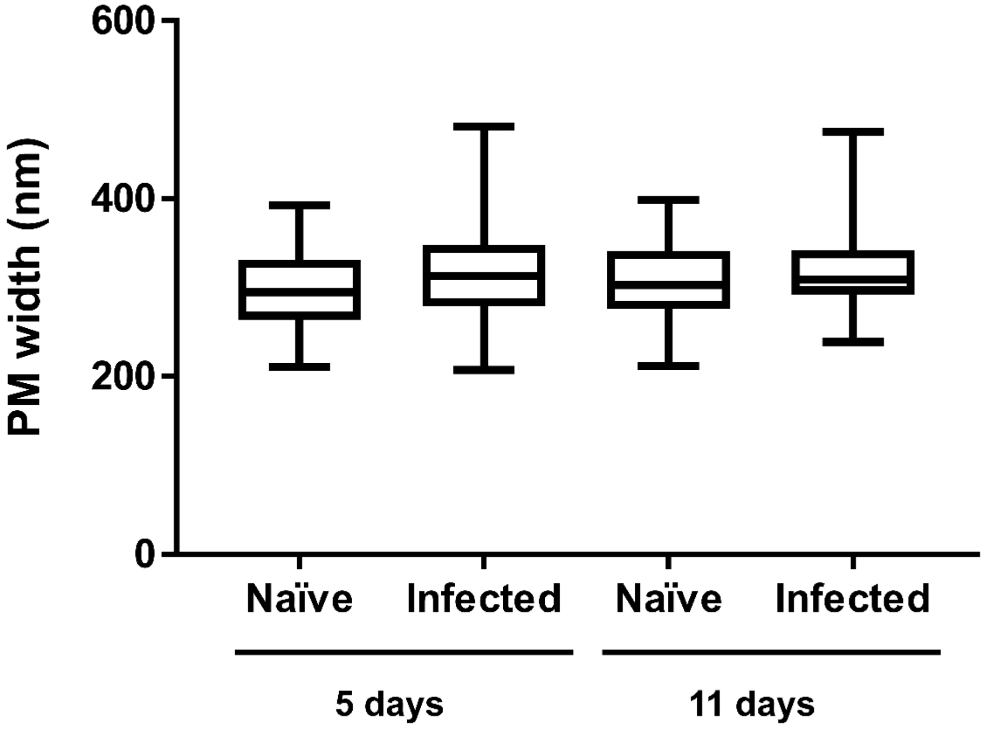

**Supplementary Fig. 7| PM width remains constant in infected and naïve flies.**
Box and whisker plots showing the distribution in width of the PM in naïve compared to infected flies, at both 5 dpi and 11 dpi. Measurements were taken between PM1 and PM3. The average width for all timepoints and conditions is ~300nm and variation is slightly increased in infected flies as these groups consider any measurements that were taken across a cyst.

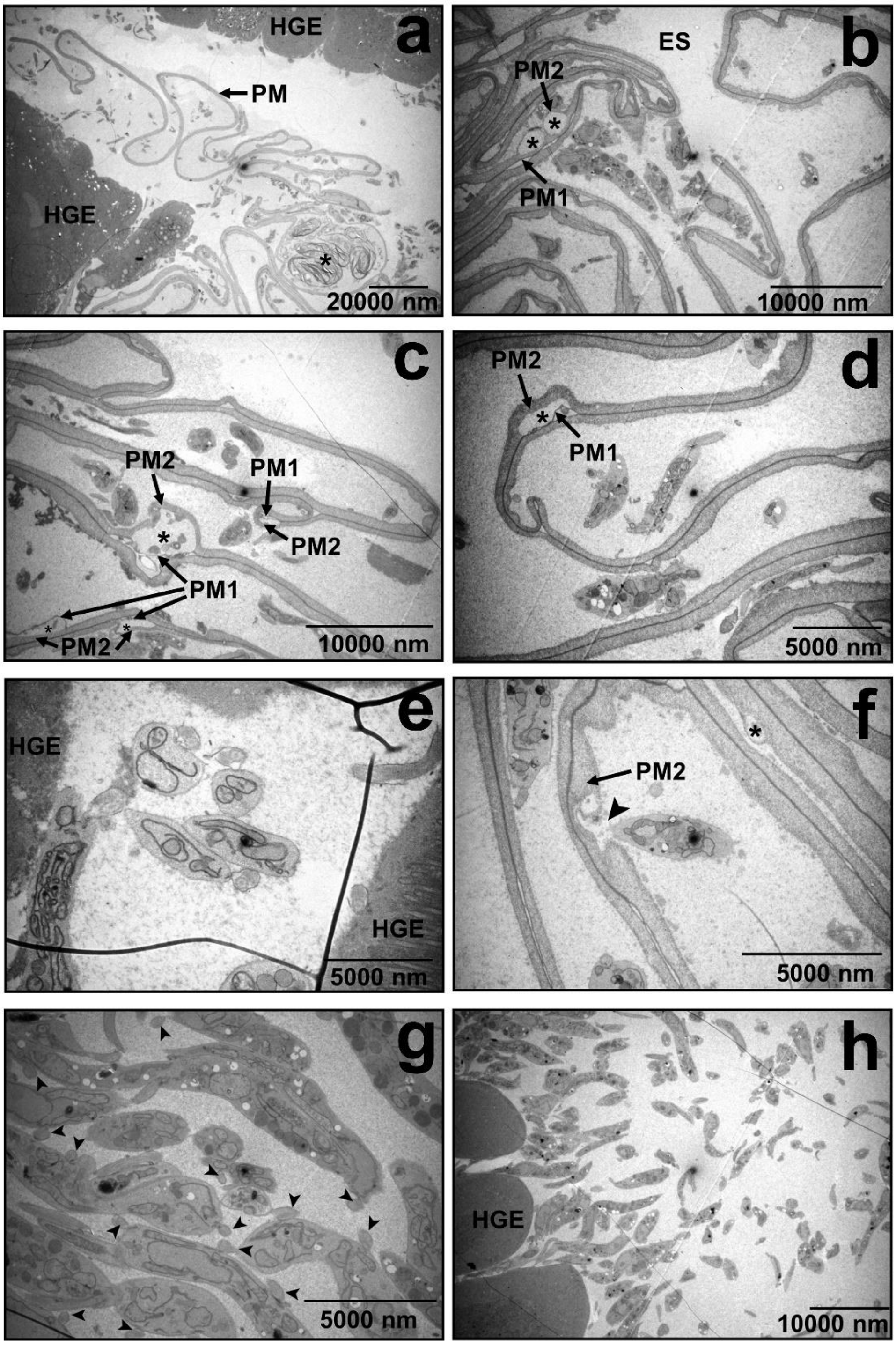

**Supplementary Fig. 8| A selection of TEM micrographs taken from the hindgut of a fly at 11 dpi. a**, An overview of the hindgut of an infected fly. The epithelial cells of the hindgut are distinctive (HGE) and the PM can be seen. In this region, material presumably from degraded and discarded PM can be seen (*) (790 x). **b-d**, Empty cysts with no visible parasites can be seen. **e**, In some instances, there appeared to be parasites that looked typical in shape to what is normally seen in the anterior midgut. **f**, A parasite close to a damaged PM2 (arrowhead). Damage can also be seen to PM2 (*) in another part of the PM. **g-h**, Trypanosomes with abnormal morphology in the ectoperitrophic space of the hindgut. Arrowheads indicate the presence of multiple flagella.

| | 5 d.p.i. | 5 d.p.i refractory | 5 d.p.i blood fed naive | | 8 d.p.i. | 8 d.p.i refractory | 8 d.p.i blood fed naive | | 11 d.p.i. | 11 d.p.i refractory | 11 d.p.i blood fed naive | Total |
|---|---|---|---|---|---|---|---|---|---|---|---|---|
| Number of midguts | 16 | 7 | 9 | | 9 | 4 | 7 | | 12 | 8 | 10 | 82 |
| Average number of grids taken per fly | 8.8 | 7 | 6.9 | | 6.1 | 8.75 | 9.8 | | 11.3 | 7 | 10.6 | |
| Average number of images per grid | 12.2 | 10.4 | 11.6 | | 12 | 14 | 7.4 | | 16 | 4.3 | 9 | |
| Number of hindguts | | | | | | | | | 4 | | | 4 |
| Average number of grids taken per fly | | | | | | | | | 5.4 | | | |
| Average number of images per grid | | | | | | | | | 11 | | | |
| Number of PVs | 7 | 4 | 5 | | | | | | 8 | 8 | 10 | 42 |
| Average number of grids taken per fly | 25.3 | 11.3 | 12.5 | | | | | | 17.2 | 6 | 10.7 | |
| Average number of images per grid | 18 | 9 | 8 | | | | | | 14 | 12 | 5 | |

**Supplementary Table 1| Total number of samples used for TEM analysis.** Numbers for each tissue represent individual flies and average numbers are from each of the different timepoints and fly conditions. Average number of grids were either sequential or taken at a distance further into the tissue.

### Supplementary Video legends

**Supplementary Video 1. Z-stack of trypanosome cyst.**

GFP expressing trypanosomes can be seen within the layers of an isolated PM from a fly at 9 dpi How many stacks. Parasite DNA is shown in magenta. Taken under 63x oil objective.

**Supplementary Video 2. Serial sections of a cyst from a fly at 11 dpi.**

474 serial sections of a trypanosome filled pocket in the PM in the anterior midgut. Sections were cut at 100nm and scanned under SBSEM

**Supplementary Video 3 Reconstruction of a cyst from a fly at 11 dpi.**

SBSEM and manual segmentation showing parasites are contained and lying parallel within PM1 and PM2 and are not seen crossing the PM.

**Supplementary Video 4. Serial sections of PV ROI1 from a fly at 5 dpi.**

458 serial sections in a region of high parasitaemia in the PV of a fly at 5 dpi Sections were cut at 40nm and scanned under SBSEM

**Supplementary Video 5. Serial sections of PV ROI2 from a fly at 5 dpi.**

457 serial sections in a second region of high parasitaemia in the PV of a fly at 5 dpi Sections were cut at 40nm and scanned under SBSEM

**Supplementary Video 6. Reconstruction of PV ROI2 from a fly at 5 dpi.**

SBSEM and manual segmentation showing the cuticular intima (yellow) of the foregut is intact, there appears to be no penetration of PV cells by trypanosomes and no indication of attachment. Four trypanosomes were also reconstructed. The scale bar is representative of the SEM image not the reconstructed image.

**Supplementary video 7. LSCM of embedded trypanosomes**

Maximum intensity projection of an isolated PM (orange) from a fly at 8 dpi. Trypanosomes (green) can be seen trapped on the ectoperitrophic side of the PM by PM material.

**Supplementary video 8. PV from a fly at 2 dpi**

PV shown in DIC with trypanosomes in green, indicated by line and circles. Trypanosomes are clearly inside the tissue and different from those swimming freely outside.

**Supplementary video 9. Annotated LSCM of naïve PV**

Z-stack of a naïve PV showing the origin of the PM (orange), F-actin structure of the PV (white) and nuclei (magenta).

**Supplementary video 10. 3D representation of naïve PV**

3D render of a PV from a naïve fly (as shown in supplementary video 9).

**Supplementary video 11. Annotated LSCM of infected PV**

Z-stack of a PV from an infected fly at 5 dpi Stains are the same as in Supplementary video 9 but with eGFP to show trypanosome distribution.

**Supplementary video 12. 3D representation of infected PV**

3D render of a PV from a fly at 5 dpi (as shown in supplementary video 11).
